# Dynamical constraints on neural population activity

**DOI:** 10.1101/2024.01.03.573543

**Authors:** Emily R. Oby, Alan D. Degenhart, Erinn M. Grigsby, Asma Motiwala, Nicole T. McClain, Patrick J. Marino, Byron M. Yu, Aaron P. Batista

## Abstract

The manner in which neural activity unfolds over time is thought to be central to sensory, motor, and cognitive functions in the brain. Network models have long posited that the brain’s computations involve time courses of activity that are shaped by the underlying network. A prediction from this view is that the activity time courses should be difficult to violate. We leveraged a brain-computer interface (BCI) to challenge monkeys to violate the naturally-occurring time courses of neural population activity that we observed in motor cortex. This included challenging animals to traverse the natural time course of neural activity in a time-reversed manner. Animals were unable to violate the natural time courses of neural activity when directly challenged to do so. These results provide empirical support for the view that activity time courses observed in the brain indeed reflect the underlying network-level computational mechanisms that they are believed to implement.

## Introduction

The time evolution of neural population activity, also referred to as neural dynamics, is believed to underlie many brain functions, including motor control^1^, sensory perception^2–4^, decision making^5–8^, timing^9,10^, and memory^11,12^, among others^13^. As examples, decisions might be formed by neural activity converging to point or line attractors^6–8;^ memories might be recovered by neural activity relaxing to a point attractor^12,14^; and arm movements might involve neural activity that exhibits rotational dynamics^1^. The similarities between the temporally structured population activity produced by network models^6–8,10,14–17^ and that produced by the brain^1,2,6–8,10^ have provided tantalizing evidence about how the brain achieves computation through dynamics^18–22^.

In network models, the time evolution of activity is specifically determined by the network’s connectivity^23^. That is, the activity of each node at a point in time is determined by the activity of every node at the previous time point and the network’s connectivity. Such neural dynamics give rise to the computation being performed by the network and are often characterized with a flow field^23^. The particulars of these flow fields reflect the specific computations performed by the network, as they arise from the connectivity of the network. Previous studies^2,4,6,7,10,16,20^ have demonstrated that activity time courses in the brain appear to follow a flow field, resembling the activity of network models. These studies suggest that the brain operates on principles like those that govern network models, but the links have not been firmly established. If the activity time courses observed in the brain indeed reflect network principles, then they should be robust and difficult to violate, since doing so could require a change to the network itself.

Here we test whether a fundamental aspect of the conceptual framework of computation through dynamics applies to biological networks of neurons by asking: to what extent is neural population activity constrained to follow specific time courses? Is it possible to produce the same population activity patterns, but in a different temporal ordering (e.g., to traverse the natural time course of neural activity in a time-reversed order)? If neural activity time courses indeed reflect the underlying network connectivity, which subserves specific computations, then the time courses should be difficult to alter.

To provide a direct test of the robustness of neural activity time courses, we employ a brain-computer interface (BCI) paradigm. In a BCI, the user is provided with moment-by-moment visual feedback of their neural activity. A BCI allows us to harness a subject’s volition to attempt to alter the neural activity they produce, and it thereby provides a powerful tool for causally probing the limits of neural function^24,25^. In prior work, we used a BCI to ask what population activity patterns an animal is able to achieve^26,27^. Here we ask whether those activity patterns can be expressed in a different temporal ordering.

We begin by identifying the naturally-occurring time courses of neural population activity in the motor cortex of Rhesus monkeys during a BCI task. We report that even during BCI control, motor cortex population activity exhibits dynamical structure, akin to what has been observed previously using arm movements^1^. In this study we interrogate that temporal structure, to see if we could lead the animals to violate it. We find that these natural activity time courses are remarkably robust. When we provided animals with visual feedback of different views of their neural activity, we observed nearly the same time courses of neural population activity regardless of the view. We then directly challenged animals to volitionally alter the time evolution of their neural population activity, including traversing the natural activity time courses in a time-reversed manner. We found that animals were not able to readily alter the time courses of their neural activity. Rather, neural activity adhered to its natural time courses, despite strong incentives to modify them. This provides evidence that the natural activity time courses reflect an underlying flow field that is difficult to violate. Our results forge a link between the activity time courses observed in a broad set of empirical studies (e.g., refs ^1,2,5–11)^ and the network-level computational mechanisms they are believed to support.

## Results

We begin with some terminology. A neural “population activity pattern” refers to the joint firing rate of a population of neurons at a moment in time. A “neural trajectory” is a time course of one neural population activity pattern, then another neural population activity pattern, and so forth, in a characteristic order on a timescale of tens of milliseconds (Fig. 1A). In this study, we asked whether the time course of neural population activity in motor cortex is easy or difficult to modify. If neural trajectories are difficult to modify it would imply that they are constrained by the underlying network. Alternatively, if neural trajectories are easy to modify, it would imply that the underlying network does not constrain the ordering in which a given set of population activity patterns can be produced (Fig. 1B, C), and thus the neural trajectories observed in the brain might be incidental.

**Fig. 1.**
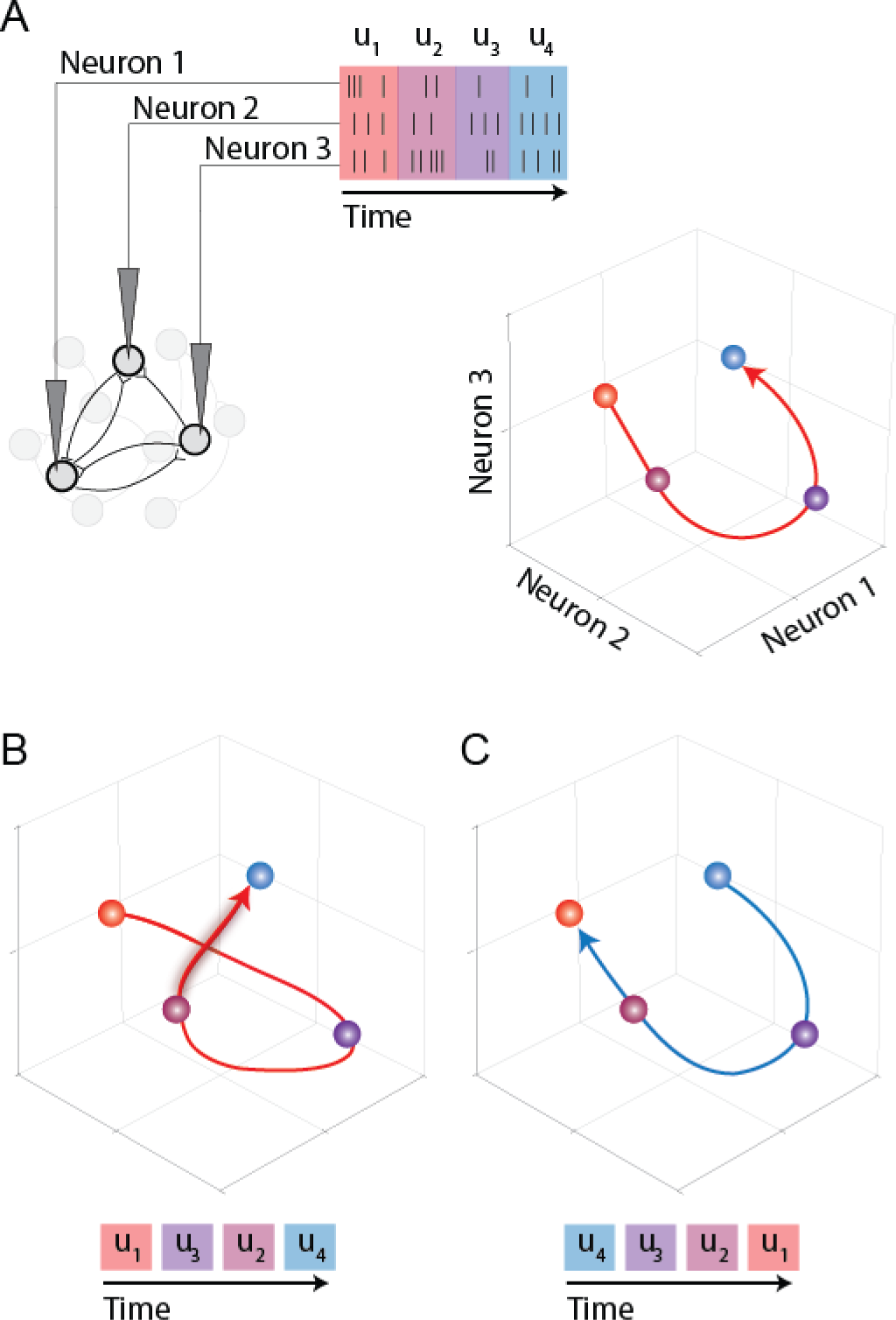
Testing the flexibility of the time courses of neural population activity. A. Activity recorded from a population of neurons is binned in time (e.g., tens of milliseconds) and represented in a population activity space, where each axis represents the firing rate of one of the recorded neurons. The time course of the population activity patterns forms a neural trajectory (red line). For illustration purposes, a three-electrode recording is shown, corresponding to a three-dimensional population activity space. In our actual experiments, ∼90 electrodes were used. B, C. The central question of this study is to ask whether neural population activity patterns can be generated in different orderings. B. An example of a different ordering of the same population activity patterns that comprise the trajectory shown in (A). In this example, the start and end points of the trajectory are the same as in (A), but the activity patterns are produced in a different order. Although it appears that the trajectory intersects with itself, in the 3D population activity space it is not intersecting. C. A depiction of the time-reversed trajectory shown in (A).

The goal of our experiments was to test the extent to which neural trajectories are flexible. To do this, we needed a way to challenge the animal to generate neural population activity patterns in a particular ordering. With a BCI, we can request population activity patterns with specific characteristics and test if the animal can produce them. To this end, we recorded the activity of a population of ∼90 neural units from each of three Rhesus monkeys, implanted with a multi-electrode array in motor cortex (Fig. 2A). The recorded neural activity was transformed into 10-dimensional latent states using a causal form of Gaussian Process Factor Analysis (GPFA^28^; see Methods). Animals then controlled a computer cursor via a BCI mapping that projected the 10D latent variables to the 2D position of the cursor (see Methods). A position mapping is a critical design choice here and is a departure from our previous work in which we mapped neural activity to cursor velocity^26,27^. By rendering neural activity as a cursor position, we provided the animal with direct visual feedback (i.e., a 2D projection) of its neural population activity unfolding over time. This makes the temporal structure in neural population activity visible to the animal. To directly challenge the animals to generate population activity patterns in a particular ordering, we designed tasks that required the BCI cursor to acquire targets or to follow paths that we specify.

**Fig. 2.**
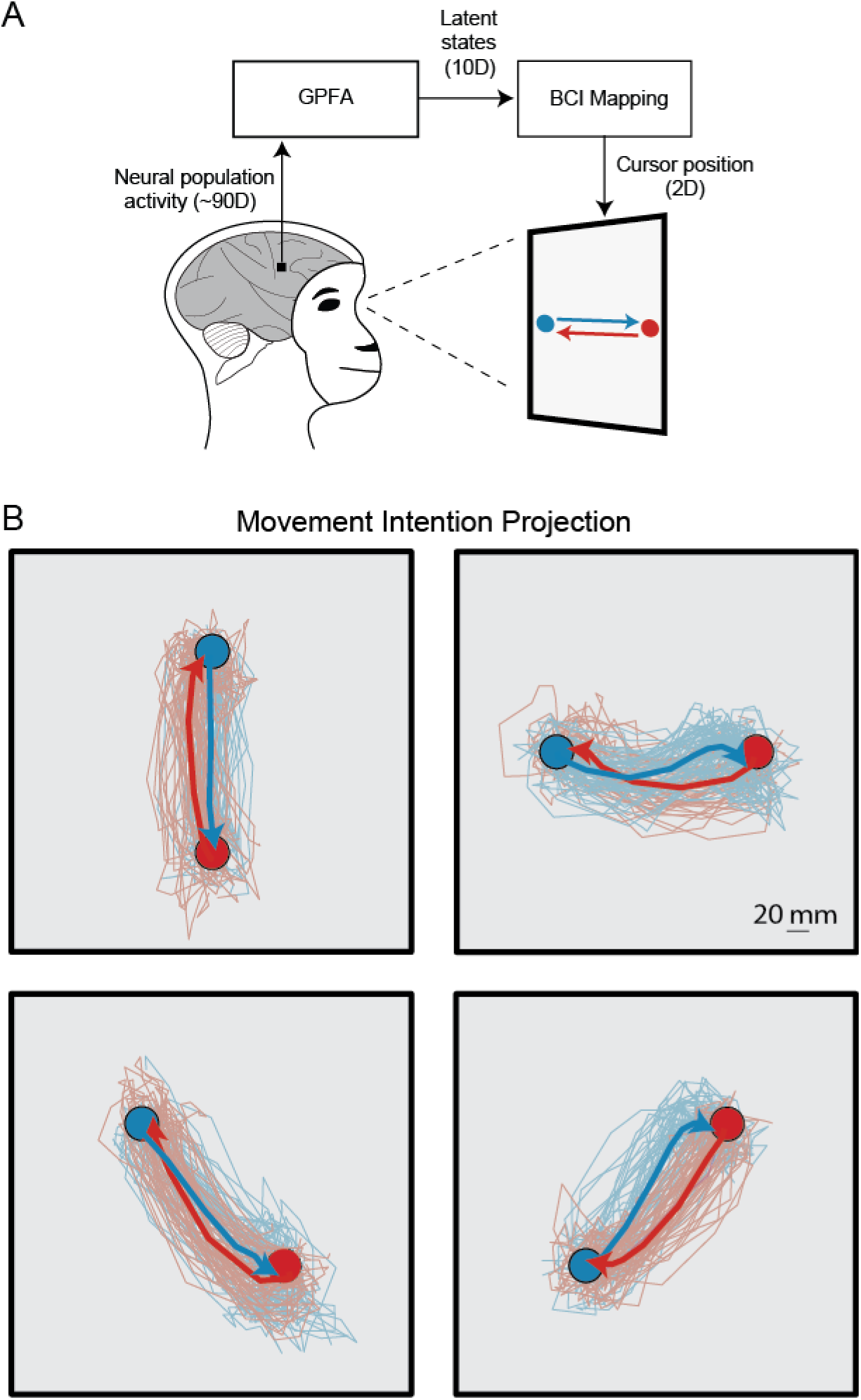
-A monkey can move the BCI cursor in any direction in the movement intention projection. A. The BCI provides the monkey with moment-by-moment visual feedback of its population activity patterns as they evolve over time. We used a causal version of Gaussian process factor analysis (GPFA)^28^ to estimate 10D latent states at each time step based on the recorded population activity (∼90D). We then provided the monkey with visual feedback of two dimensions of its latent states (defined by a BCI mapping, see Methods), which determine the moment-by-moment 2D position of the BCI cursor. The monkey moved the cursor from target A to target B (blue) and from target B to target A (red). B. Cursor trajectories for the four possible target pairs while using the movement intention (MoveInt) BCI mapping. For visualization, trajectories are colored by the start target (e.g., red trajectories indicate movements originating at the red target and moving to the blue target). Thin traces represent individual trial trajectories. Thick traces represent trial-averaged trajectories. Note the high degree of overlap between the red trajectories and blue trajectories for each target pair.

A key experimental design choice is to specify which 2D projection of its 10D neural trajectories we show to the animal. Each experiment began with a BCI mapping that was intuitive to use and captured the monkey’s movement intention, i.e., the “movement intention” (MoveInt) mapping (see Methods). With the MoveInt mapping, the animals could move the cursor flexibly throughout the workspace. Animals performed a two-target BCI task in which they moved the cursor between a pair of diametrically opposed targets. In the MoveInt projection, the cursor trajectories from target A to target B were highly overlapping with the cursor trajectories from target B to target A (Fig. 2B). The overlapping cursor trajectories might lead one to believe that, from a given position in 10D space, it would be possible to move in multiple different directions. That is, this projection makes it appear that the time courses of population activity patterns are flexible.

The MoveInt BCI mapping is just one 2D projection of the 10D space of neural population activity. Although the cursor trajectories between a given target pair overlap in the MoveInt projection, the neural trajectories are defined in a 10D space and need not overlap in all dimensions. In fact, when we view other projections of the 10D space, the temporal structure of the neural trajectories differed in a direction-dependent way (Fig. 3A). In some 2D projections, we found that the neural trajectories when moving the cursor from target A to target B were distinct from the neural trajectories when moving the cursor from target B to target A (Fig. 3B). These direction-dependent paths show that the monkey did not simply produce the same population activity patterns in a different ordering to move back and forth between the targets. Rather, neural activity followed different paths in 10D space, that projected onto overlapping paths in the MoveInt projection. We identified a 2D projection that maximized the separation between the A-to-B and B-to-A trajectories, i.e., the “separation-maximizing” (SepMax) projection (Fig. 3C; See Methods). The neural trajectories that are overlapping in the MoveInt projection are clearly separable in the SepMax projection (Fig. 3D). Neural activity in the SepMax projection resembles that seen during studies of motor cortex during reaching^1^, and is suggestive of underlying network constraints that govern the time courses of neural activity in the population activity space.

**Fig. 3.**
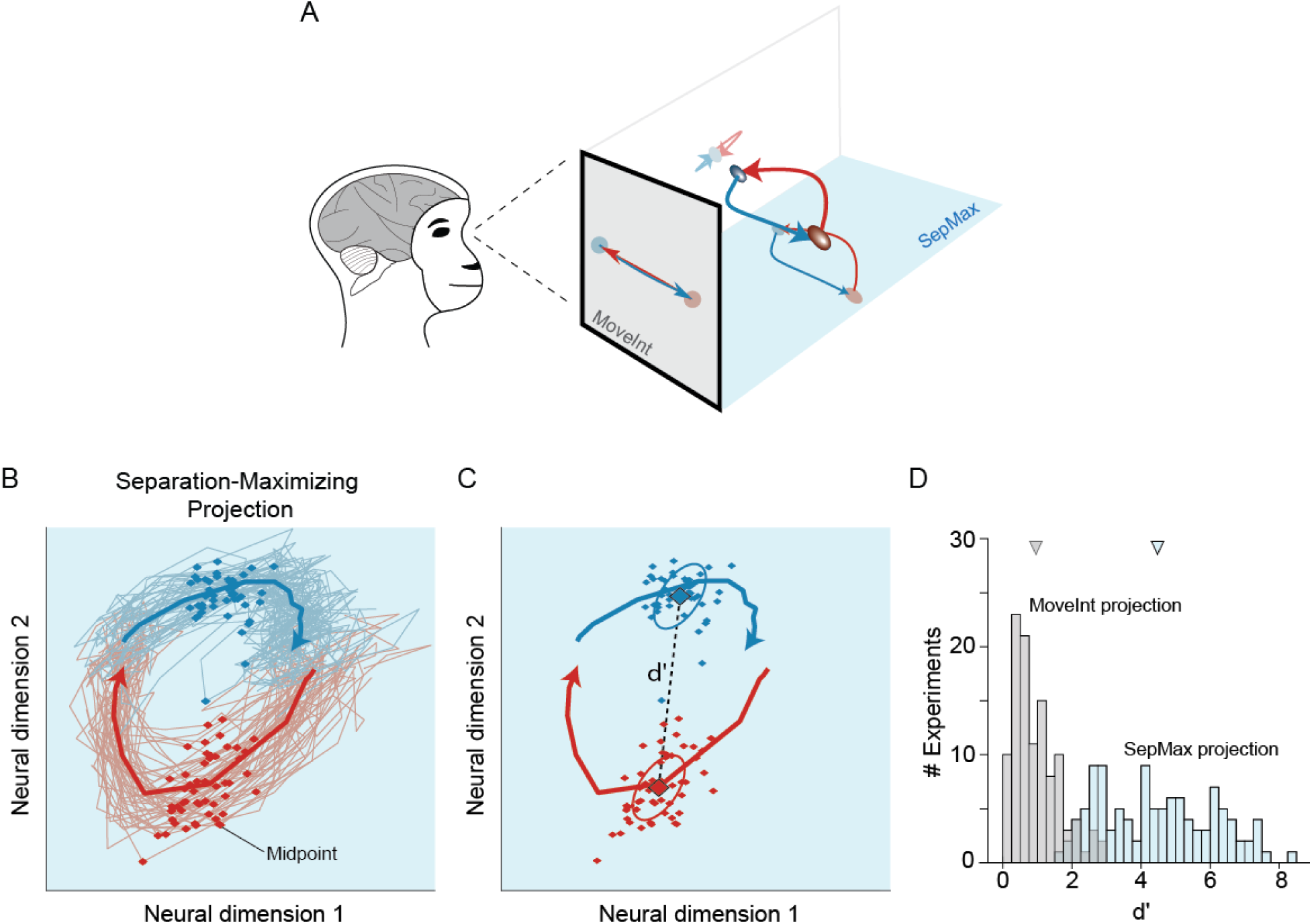
–Neural trajectories follow direction-dependent paths in the 10D population space. A. The MoveInt mapping provides the monkey feedback of one particular 2D projection of the neural trajectories. In the MoveInt projection (indicated by the gray background), the A-to-B (blue) and B-to-A (red) trajectories are overlapping. However, when we examine the 10D space in which the neural trajectories reside, we find that the A-to-B (blue) trajectories are distinct from the B-to-A (red) trajectories. B. An example 2D projection of the neural trajectories in which the A-to-B (blue) trajectories are distinct from the B-to-A (red) trajectories. The separation-maximizing projection is indicated by the light blue background. Thin traces show trajectories for individual trials. Diamonds indicate the midpoint (see Methods) for each trial. Thick traces represent trial-averaged trajectories. C. We quantify the separation between red and blue trajectories in 10D space as the discriminability (*d’*) of the blue and red midpoints. Computing *d’* involves the separation of the means (dashed line) and the covariances (ellipses) of the trial-to-trial scatter (see Methods). D. Across all experiments the neural trajectories are substantially more separated in the SepMax projection (*d’* = 4.5 ± 1.6; mean ± std) than they are in the MoveInt projection (*d’* = 0.9 ± 0.6; mean ± std) (t-test, p<10^-10^).

Once we identified this temporal structure in motor cortex activity during BCI control, we tested the flexibility of the time courses of neural population activity with three increasingly stringent experimental manipulations. First, we gave the animal visual feedback of the dimensions where temporal structure was evident. Next, we asked if the animal could produce population activity patterns in a time-reversed neural trajectory. Finally, we directly challenged the animal to follow a prescribed path through a 2D projection of the 10D neural population space. We present these experiments below.

In our first test of the flexibility of the time course of neural activity, rather than providing the monkey visual feedback of its neural trajectories in the MoveInt projection (Fig. 4A, also Figs. 2 and 3), we provided visual feedback in the SepMax projection (Fig. 4B). The animal performed the same two-target task, moving the BCI cursor between targets A and B, but now with feedback of its neural activity in a projection in which we had observed directionally-dependent curvature of the neural trajectories (Extended Data Fig. 1). We wondered if, given this feedback, the direction-dependent paths would persist (Fig. 4C, possibility 1) or if the monkeys would instinctively straighten out the trajectories (Fig. 4C, possibility 2). In behavioral studies, both humans^29–33^ and animals^34,35^ tend to adjust their movements to straighten out visually curved cursor trajectories. We leveraged this behavioral tendency to test how flexible neural activity time courses are by showing animals their curved neural trajectories. If the monkeys straightened out their cursor trajectories, that would show that the time courses of neural activity are flexible in the 10D neural population space. However, if they do not straighten out their trajectories, that would be consistent with the possibility that activity time courses are obligatory.

**Fig. 4.**
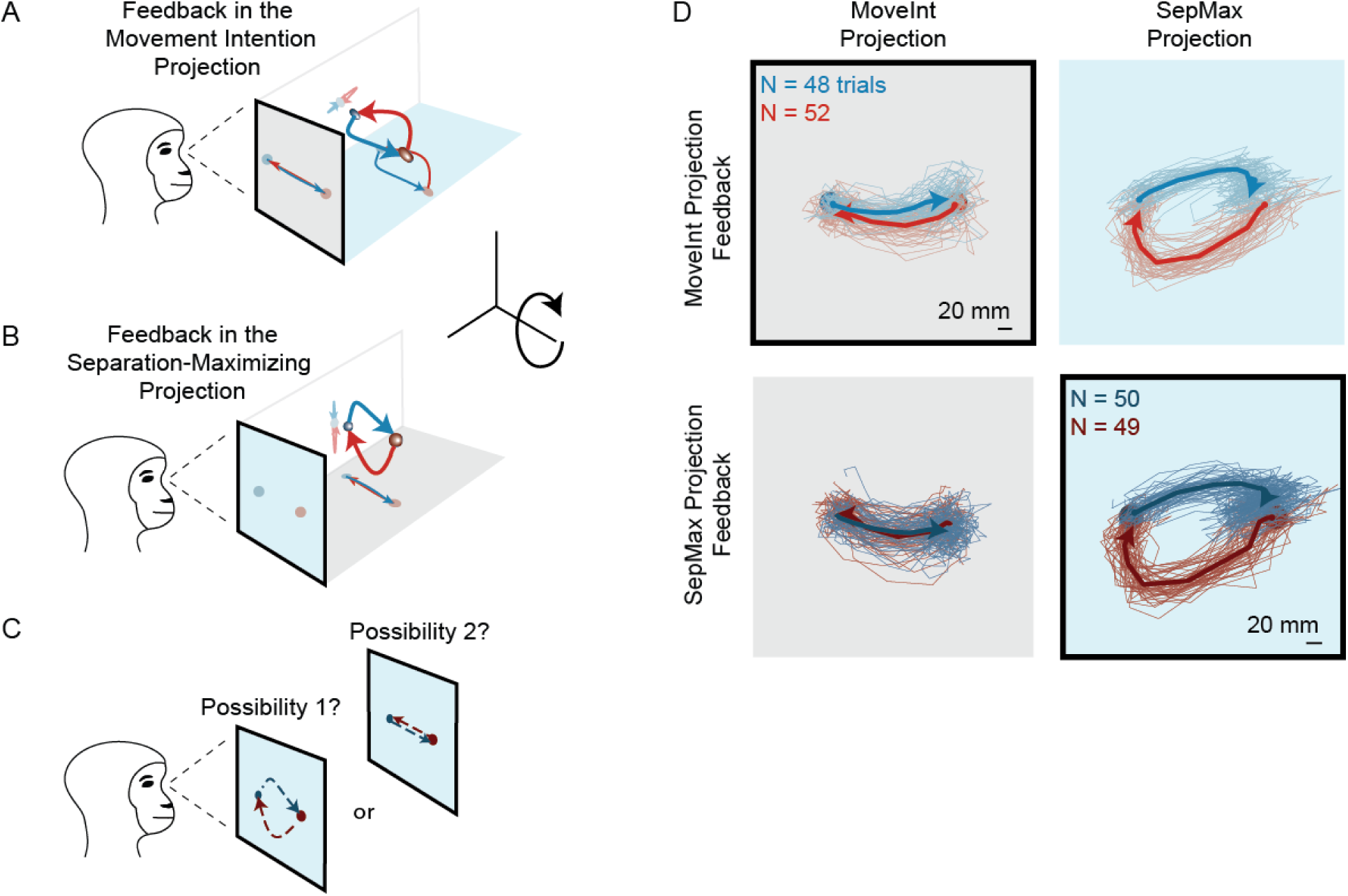
– Direction-dependent paths of neural trajectories persist even when BCI feedback is given in the separation-maximizing projection. The BCI paradigm allows us to choose which 2D projection of the neural trajectories we provide the monkey as visual feedback. Rather than show the monkey the A. MoveInt projection (gray background), we rotate the population activity space relative to the monitor to show him the B. SepMax projection (light blue background). C. Under the SepMax projection, we tested if the direction-dependent paths observed during the MoveInt block would persist (possibility 1) or if the monkey would straighten out its cursor trajectories (possibility 2). D. Neural trajectories are similar whether the MoveInt or SepMax projection is used for feedback. When the monkey received visual feedback of the MoveInt projection (top row), the A-to-B trajectories (blue) overlapped with the B-to-A trajectories (red) in the MoveInt projection (top left; gray background. Black outline indicates the projection which the monkey views on the monitor. Those same trajectories are distinct in the SepMax projection (top right; light blue background). When the monkey received visual feedback of the SepMax projection (bottom row), the trajectories continued to follow direction-dependent paths (bottom right; light blue background with black outline). Those same trajectories overlap in the MoveInt projection (bottom left; gray background).

When animals were given visual feedback in the SepMax projection, their cursor trajectories continued to show strong persistence of direction-dependent paths (Fig. 4D, bottom right panel), consistent with possibility 1 (in Fig 4C). In fact, this manipulation resulted in little change to the neural trajectories: the trajectories in the SepMax projection when it was being viewed by the animal (Fig. 4D, bottom right panel) were similar to the trajectories in the SepMax projection when the animal was receiving BCI feedback of the MoveInt projection and the SepMax projection was unseen by the animal (Fig 4D, top right panel). This was also true for neural trajectories in the MoveInt projection (Fig. 4D, left column) – trajectories were similar in the MoveInt whether or not the animal was observing that projection. Providing animals feedback of their neural trajectories in the SepMax view did not lead the animal to straighten out the trajectories in that projection, as we might have expected if the animal had volitional control over its trajectories in that projection. This finding was highly consistent across sessions (Supp. Fig. 2).

We used a “flow field” analysis to quantify the characteristic, direction-dependent paths of the neural activity time courses. Computed from the data, a flow field describes how neural trajectories unfold from any given location in neural population space. We identified the flow fields for the MoveInt and SepMax projections for both the MoveInt and SepMax feedback conditions (Fig. 5A; see Methods and Extended Data Fig. 3). We observed that the flow field in the SepMax projection was similar, whether or not it was the projection viewed by the animal (Fig. 5A, blue arrow). By comparison, the flow fields in the feedback projections (i.e., MoveInt projection with MoveInt feedback versus SepMax projection with SepMax feedback) were quite different (Fig. 5A, black arrow). To quantify these observations, we calculated the mean squared error between the flow fields (see Methods; Fig. 5B). The difference between the flow fields was smaller for a given 2D projection in different feedback conditions than between different 2D projections (Fig. 5B; the points lie below the diagonal). The robustness of the flow fields under different visual feedback projections supports the hypothesis that neural population activity is constrained to follow specific time courses.

**Fig 5.**
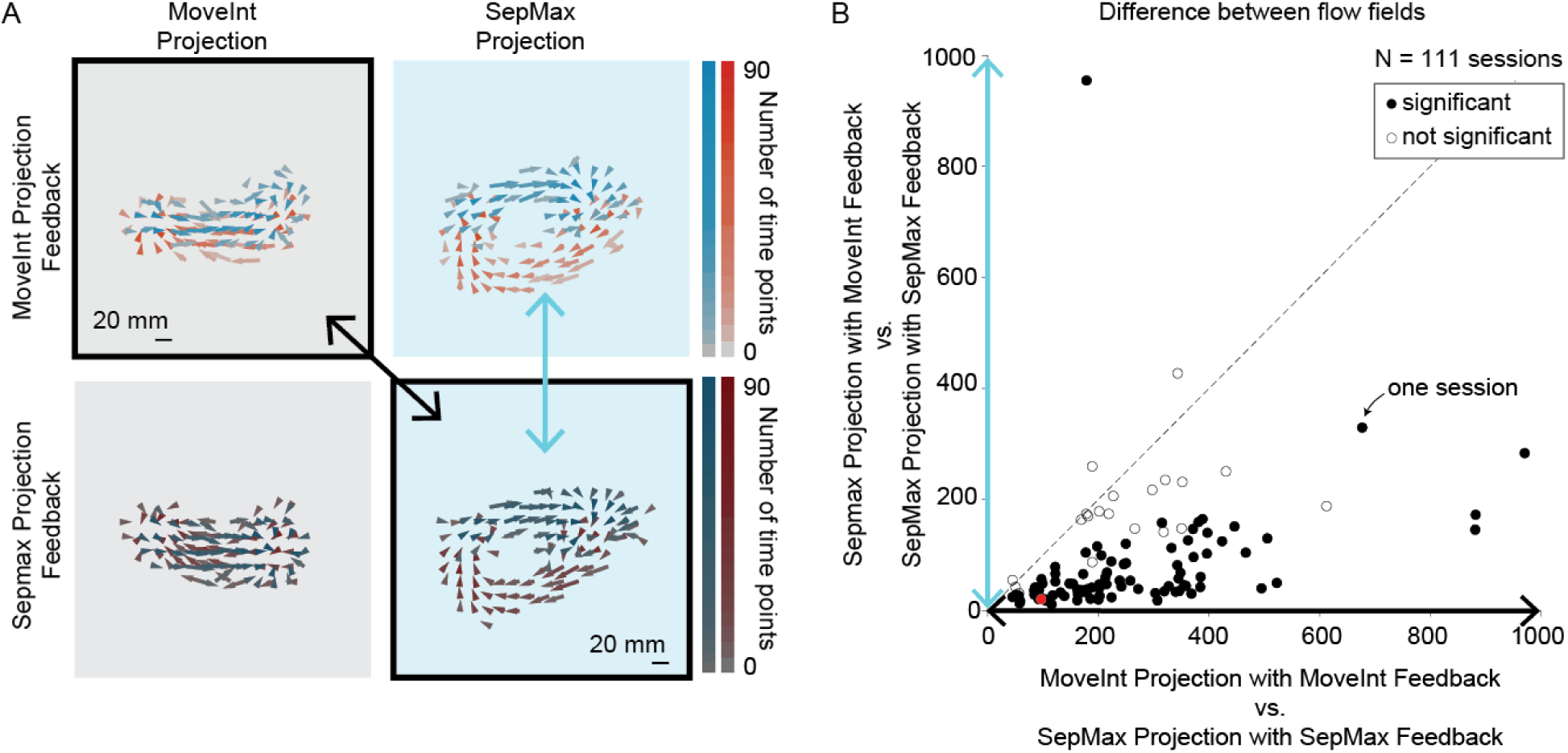
. Neural trajectories are robust to which projection provides the visual feedback. A. We can compute a flow field to summarize the cursor trajectories (see Methods). Each panel corresponds to a panel in Fig. 4D. Length and direction of arrows indicate the average, observed cursor velocity as a function of position in the corresponding 2D space. Arrows are colored by the direction of the cursor movement, and the color saturation indicates the number of data points that contribute to the average. The flow fields in the SepMax projection when seen (black outline) and unseen (no outline) by the monkey were similar (comparison noted by large light blue arrow). In contrast, the flow field in the SepMax projection differed from the flow field in MoveInt projection (comparison noted by large black arrow). B. We quantified the similarities between the flow fields by calculating the mean squared difference of the corresponding flow field vectors (see Methods). The flow fields in the SepMax projection were similar regardless of whether they were seen or unseen by the monkey (vertical axis, cf. large light blue arrow in panel A). For reference, we considered a case where the flow fields were different, namely for the two projections that we used to provide visual feedback to the monkey (horizontal axis, cf. large black arrow in panel A). Filled symbols indicate sessions where the flow field difference in the SepMax projection regardless of feedback projection (vertical axis) was smaller than the flow field difference between the different feedback projections (horizontal axis, Wilcoxon rank sum, p<0.05, 90/111 sessions). Red dot indicates the example session shown in (A).

We wondered whether these temporal constraints were limited to specific subspaces, or whether the time courses of neural activity were constrained in dimensions other than the SepMax projection. We found that the time courses of neural activity are highly consistent in all dimensions, regardless of the view provided to the animal (Extended Data Fig. 3). This indicates that the robustness of the temporal structure captured by the SepMax projection is not special. In all 10 dimensions of the neural population space, the animals cannot readily modify the characteristic activity time courses.

The persistence of the characteristic paths of the neural trajectories enables an additional demonstration of the robustness of the temporal structure. In a set of control experiments, we provide the monkey feedback of its neural trajectories in the SepMax projection and the “reflected” SepMax projection in separate blocks of trials. We leveraged the fact that we can reflect the orientation of the identified SepMax projection about the target axis (Extended Data Fig. 4A). This manipulation changes the direction-dependent trajectory curvature. For example, if the cursor trajectories from target A to B curved upwards in the SepMax projection, the trajectories would curve downwards in the reflected SepMax projection. With feedback in the reflected SepMax projection, one possibility is that the trajectories might continue to exhibit the same direction-dependent trajectory curvature that we observed before the reflection (Extended Data Fig. 4B, possibility 1). This possibility would demonstrate flexibility in the activity time courses and would indicate that the animal simply had a preference for generating population activity patterns along particular paths. Another possibility is that the cursor trajectories might show the corresponding reflection of the characteristic paths in the reflected SepMax projection (Extended Data Fig. 4B, possibility 2). This possibility would be consistent with rigid temporal structure. Remarkably, when we provided the monkey feedback of its neural trajectories in the reflected SepMax projection, the cursor trajectories were also reflected (Extended Data Fig. 4C, D). This observation further strengthens our finding of a robust temporal structure that likely reflects network constraints.

Taking stock, we first observed temporal structure in dimensions of neural population activity that were unseen by the animal during BCI control (Fig. 3). When we provided visual feedback of those dimensions to the monkey, the characteristic time courses of neural activity persisted (Fig. 4, Extended Data Fig. 4). In our next test, we sought a more direct test of the flexibility of neural trajectories. We asked if the animal could produce previously observed population activity patterns in a time-reversed ordering. To do this, we used the empirically-derived flow fields (Fig. 5) to describe the characteristic paths of the neural trajectories in the SepMax projection (Fig. 6A). Then, we challenged the animal to move the BCI cursor against the flow field (Fig. 6B, large red arrow). We presented a target along one of the paths of the flow field (Fig. 6C). This “intermediate target” (black target) was naturally acquired from target A (i.e., the blue path from target A to the intermediate target follows the flow field) but moving the cursor from target B (i.e., red target) to the intermediate target challenged the monkey to order its population activity patterns into a neural trajectory that moved against the flow field.

**Fig. 6.**
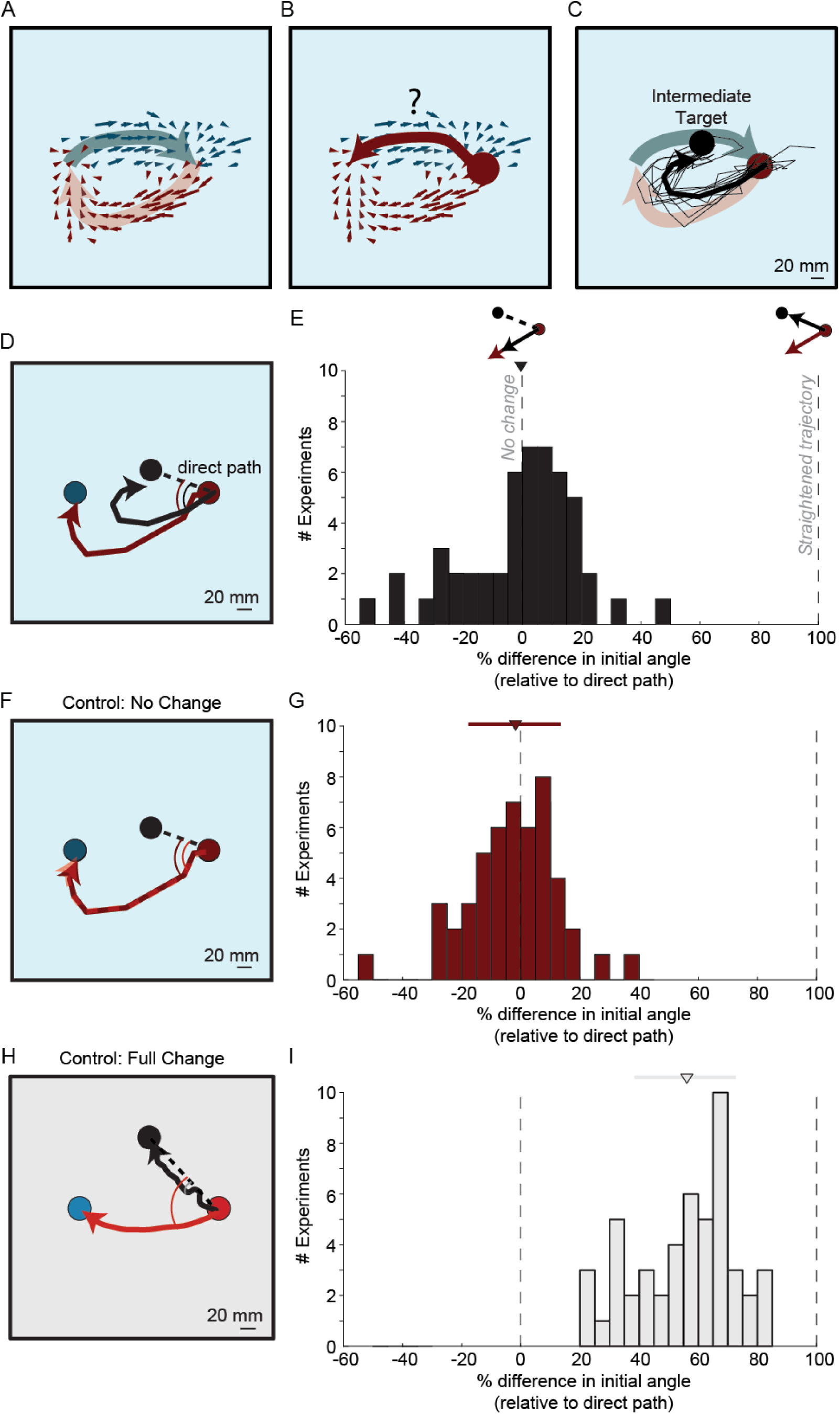
Challenging the monkey to violate the flow field. A. Flow fields (small arrows) characterize the distinct paths (represented by the large arrows) as the monkey moves the cursor from A-to-B (blue) and from B-to-A (red). Same conventions as in Fig. 5. B. We asked if the monkey could produce a time-reversed neural trajectory. For example, starting at target B (red circle) can the monkey produce a trajectory (red arrow) that moves against the flow field (blue arrows)? C. To challenge the monkeys to move against the flow field, we placed an “intermediate target” (black circle) along the path of the flow field. Black lines show single-trial cursor trajectories (thin lines) and trial-averaged cursor trajectory (thick line) to the intermediate target. From target B (red circle), the cursor initially follows the flow (red arrow) before hooking to acquire the intermediate target (black circle). D. Quantifying the animal’s ability to move the cursor against the flow field. We measure the initial angle between the cursor trajectory to the intermediate target (black line; same as in panel C) relative to the direct path to the intermediate target (dashed black line) and then averaged across trials. For comparison, we measure the initial angle between the cursor trajectory to the blue target in the two-target task (red line; same as in Figs. 3C and 4D, but using only the early half of the trials, see Methods) and the direct path to the intermediate target and then averaged across trials. E. Distribution of the percent difference of the two initial angles illustrated in panel D across experiments. Zero percent difference (vertical dashed line, left) indicates that the cursor moves along the flow field before acquiring the intermediate target. One hundred percent difference (vertical dashed line, right) indicates that the cursor was able to move straight to the intermediate target (against the flow field). F. For the no change control, we compute the initial angles between the early (red line, same as in panel D) and late (dashed red line) two-target trials relative to the direct path to the intermediate target (dashed black line). G. The distribution of the difference in initial angles across experiments for the no change control. H. For the full change control, we use cursor trajectories under the MoveInt projection (cf. Fig. 2B), where the animal has flexible control of the cursor and can easily move from a start target (red circle) to targets (blue and black circles) that are in approximately the same position as the blue target and intermediate target in the SepMax projection. We compute the initial angles between each of these trajectories (red line and black line) relative to the direct path to the black target (dashed black line). I. The distribution of the difference in initial angles across experiments for the full change control.

Importantly, the animal was not asked to generate *new* population activity patterns, but rather to generate previously-observed patterns in a new ordering. When the monkeys moved the cursor from target B to the intermediate target, they did not generate the time-reversed neural trajectory to move directly to the intermediate target against the flow field. Rather, they acquired the target by initially following the path of the flow field, and then hooking back toward the intermediate target (Fig. 6C, black traces). To succeed at the intermediate target task, the animals did not violate the flow field, but instead appear, at least initially, to follow the flow field.

We sought to understand to what extent the trajectories during the intermediate target task follow or violate the flow field defined by the two-target trajectories (Fig. 6A). We asked if the initial part of the cursor trajectories in the intermediate target task (Fig. 6D, black arrow) were more similar to the cursor trajectories in the two target task (Fig. 6D, red arrow) or the time-reversed, direct path (Fig. 6D, black dashed line). We defined the initial angle of the cursor trajectories of both tasks with respect to the time-reversed direct path. Then we calculated the difference in the initial angle between the two-target trajectories and the intermediate target trajectories (Fig. 6E; see Methods). Across experiments there was no systematic difference in initial angle for the intermediate target trajectories relative to the two-target trajectories (−0.56 ± 19.61 %; mean ± std). To interpret this change in initial angle, we needed to compute two reference distributions corresponding to a “no change” condition and a “full change” condition. In the no change condition (Fig. 6F, G), we computed the difference in the initial angle for two partitions of the two-target trials. In this case, there is no experimental reason to expect a difference in initial angle. By contrast, the full change condition (Fig. 6H, I) captures the amount of change we observe when the animal has flexible control of the cursor. The change in initial angle in the intermediate target task (Fig. 6E) is not statistically different from the “no change” condition (Fig. 6G; t-test, p=0.835) but is statistically different from the “full change” condition (Fig. 6I, t-test, p <<0.001). This indicates that activity time courses are not readily modified.

With the experimental manipulations described thus far, we observed only minimal modification of the cursor trajectories. However, those tasks did not *require* that the animals generate the time-reversed neural trajectory for success. The possibility remains that the animals *can* modify their neural trajectories, but they were not sufficiently incentivized to do so. In our third test, we imposed a visual boundary around the time-reversed trajectory (Fig. 7A). This “instructed path task” constrains the path that neural activity can take to the intermediate target. Now, to succeed at the task the animal had to move the cursor to the intermediate target without exiting the boundary^36^. We gradually reduced the size of the boundary to approach the direct path to the intermediate target, which represents the time-reversed neural trajectory (Extended Data Fig. 5; see Methods). In this way, the instructed path task directly challenged the animal to modify its neural trajectories. Importantly, the allowable path includes the population activity patterns that comprised the previously-observed neural trajectories. This means that we know that the animal can produce each of the required population activity patterns, but this task challenges the animal to produce them in a different ordering. If the animal can produce the time-reversed neural trajectory, the cursor would move directly to the intermediate target. If the animal cannot alter the time course of its population activity patterns, then it would not be able to succeed at the task.

**Fig. 7.**
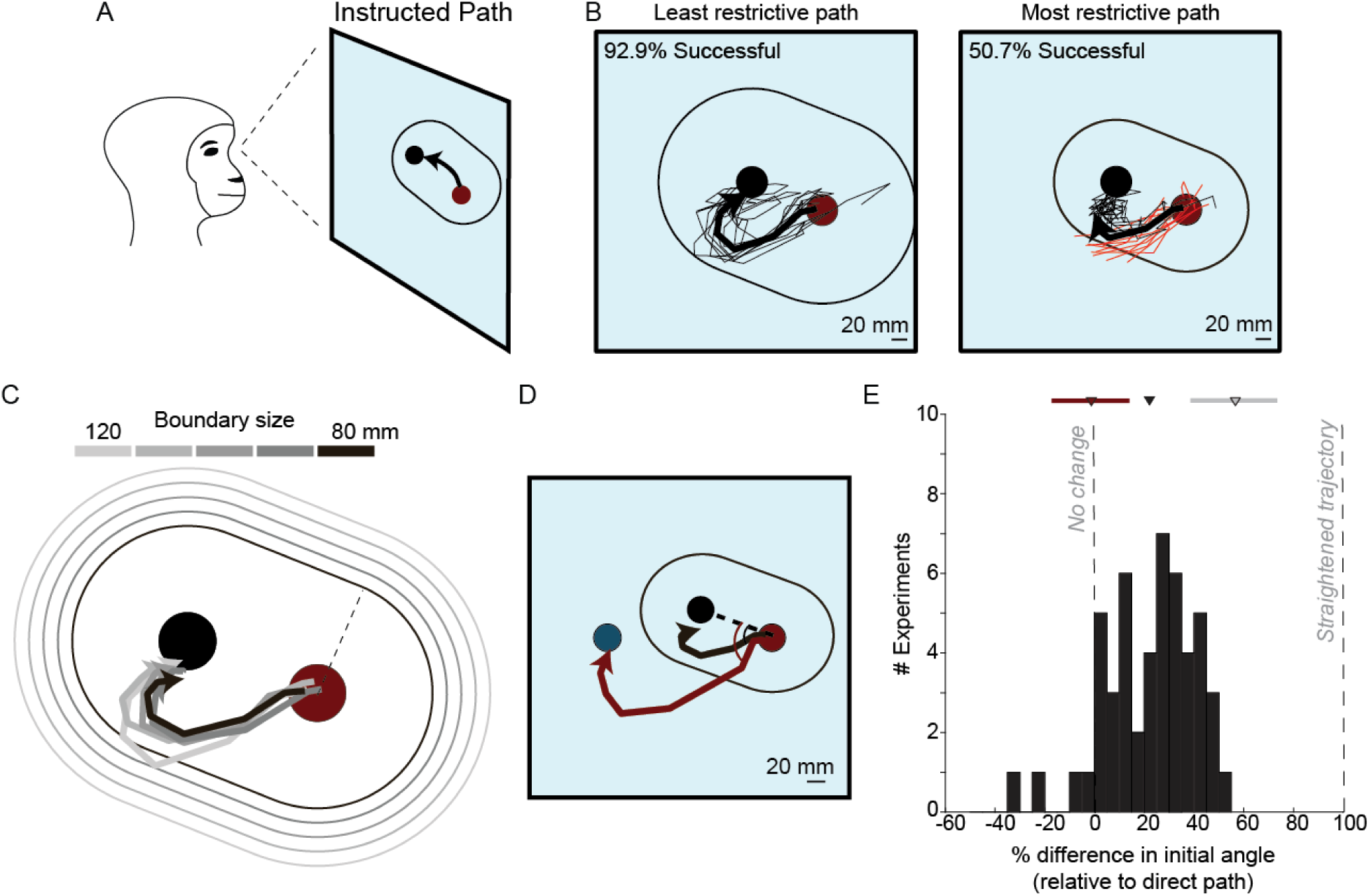
– Monkeys cannot generate time-reversed neural trajectories. A. The monkeys performed an “instructed path task” in which they had to move the cursor from the start target (red circle) to the intermediate target (black circle) without exiting a visual boundary (oval outline). B. To encourage the monkeys to modify their trajectories, we incrementally reduced the size of the boundary so that an inability to alter their trajectories would eventually lead to a failure at the task. Cursor trajectories for the least and most restrictive path for an example session. The least restrictive path (left) minimally affected the cursor trajectories, relative to the unconstrained trajectories. With the most restrictive path, the animal only succeeded approximately half the time (right). Successful trials shown as thin black lines. Failed trials shown as thin red lines. The thick black line shows the average of all trials, regardless of success, for that boundary size. C. The five boundary sizes (ovals outlines in different shades of gray; size refers to the distance of the boundary from the target indicated by the dashed line) and trial-averaged successful cursor trajectories for each boundary size (line in corresponding shade of gray) for the example session. D. Comparison of the initial angle of the most constrained trials (trial-averaged trajectory of all initiated trials regardless of eventual success, black line) to the initial angle of the two-target trials (trial-averaged trajectory of all initiated trials regardless of eventual success, red line; same as in Figs. 6D and 6F). We calculated the initial angles relative to the direct path (black dashed line) from the start target (red circle) to the intermediate target (black circle). E. Distribution of the percent difference in initial angle across experiments. The mean is indicated by the black triangle. For reference, the no-change distribution (dark red triangle and line; mean ± standard deviation) and the full change distribution (gray triangle and gray line; mean ± standard deviation) from Fig. 5 are shown here.

Even in this task with strong incentive to produce time-reversed trajectories, animals only minimally modified their trajectories as we reduced the size of the boundary. As the boundary became more restrictive, instead of modifying their trajectories, they began to fail at the task (Fig. 7B, compare left and right panels). Across all sessions, we did not observe cursor trajectories that went directly to the intermediate target, even though the animal was previously able to generate each particular population activity pattern along that path. Instead, the trajectories continued to show the hook-like bowing feature along the natural direction of flow (Fig. 7B, C; see Extended Data Fig. 6 for more example sessions) like that we observed in the intermediate-target experiment (Fig. 6C). This shows that even when given a strong incentive to reverse the ordering of neural population activity, the monkeys were unable to do so.

To quantify the flexibility of neural trajectories, we again used the change in initial angle metric. We examined the initial angle of the trajectories in the presence of the most restrictive boundary in comparison to the two-target trials (Fig. 7D). Across experiments there was a small (22 ± 18 %; mean ± std) change in the initial angle of the trajectories compared to the “no change” condition (Fig. 7E; t-test, p<10^-5^). As in the intermediate target task (Fig. 6D, E), during the instructed path task the monkeys produced trajectories that were more similar to the two-target trajectories than the direct path (Fig. 7D, E). That is, the animals produced cursor trajectories that followed the flow field, despite the presence of the boundary compelling them to alter their trajectories. This lack of flexibility demonstrates that strong constraints exist on the time courses of neural activity.

Finally, we considered whether the animals simply did not understand the instructed path. Arguing against this interpretation, the animals performed better at the instructed path than predicted from applying the boundaries to the unconstrained trials (Extended Data Fig. 5B). This shows that they were attempting to respond to the presence of the boundary. Even with these small changes in behavior, the trajectories exhibited in the instructed path task continued to resemble the direction-dependent curvature of the unconstrained trajectories.

## Discussion

Here we assessed the flexibility of the dynamical structure present in neural population activity. We first observed the naturally-occurring time courses of neural population activity in motor cortex during brain-computer interface (BCI) control. We found that these time courses exhibited rich temporal structure while monkeys used a BCI, reminiscent of what is seen in the motor cortex during reaching^1^ and elsewhere in the cerebral cortex during other behavioral tasks^2,5–7,9,20^. This temporal structure persisted when animals were given visual feedback of their neural population activity in projections that make the temporal structure most evident. When challenged to generate time-reversed versions of naturally-occurring time courses, we found that animals were unable to do so, even when strongly incentivized. These results indicate that the temporal structure of neural population activity that we, and many others (e.g., refs ^1,2,5–7,9–11)^, have observed reflects underlying network constraints that are difficult to violate.

Why should neural activity exhibit temporal structure? Temporal structure, often referred to as neural population dynamics, is taken as a signature of the computation carried out by the network of neurons. This understanding of temporal structure in neural population activity originated in neural network modeling studies^14,15,18^, in which the time-evolution of activity within a network is shaped by the connectivity of the network, and embodies the computation being performed. Once provided with an input, the network evolves from its initial state to a final activity state that provides the output of a computation (e.g., a commitment to a decision or a plan for a specific movement). Examples of temporal structures include the flow of activity toward a point attractor, line attractor, or a stable limit cycle. Empirical studies have demonstrated that such temporal structures may underlie arm movements^1^, olfaction^2^, sensory perception^3,4^, decision making^5–8^, timing^9,10^, and more. Our work provides causal evidence that the activity time courses observed in the brain in those studies are not arbitrary, but instead likely reflect the underlying network connectivity which gives rise to the relevant computations.

In network models, activity is a function of both the connectivity of the network and the inputs to the network. Thus, to observe flexible temporal structure in the activity of a network would require altering the network’s connectivity, or altering its inputs. Our findings of inflexible temporal structure lead us to conclude that neither approach was available to the monkeys, at least on the brief timescales of individual days in which we conducted our experiments. That is, they were unable to either adjust M1’s local connectivity or adjust the high-level commands sent to M1 in a manner that would allow them to alter the natural time course of their neural activity. M1’s population dynamics have been shown to depend on the inputs to M1, for example from the thalamus and other brain areas^37–40^. Our findings suggest that there are limits to the extent to which inputs can change population dynamics. It will be important for future studies to distinguish the relative contributions of local connectivity versus inputs to the shaping of cortical population dynamics.

A reasonable concern is that the animals were not sufficiently motivated to modify their trajectories, or that they did not understand the task. We consider these explanations unlikely because these well-trained and highly-motivated animals persisted in the face of challenging tasks, even when they involved blocks of hundreds of trials with low success rates. Although the animals did show some modest improvements in task success (Extended Data Fig. 5), the improvements did not reflect flexible control of their activity time courses (Fig. 7). These small improvements indicate that the animals understood what was asked of them, but were nevertheless unable to alter the temporal structure of their neural population activity.

The temporal structure we find here (cf. Fig. 3) is reminiscent of the rotational dynamics shown by Churchland and colleagues^1^ during arm movements. Subsequent studies showed that reversing the kinematics of the hands is not sufficient to reverse the direction of those rotational dynamics^41^. Here the temporal structure is evident without arm movements. This indicates that the temporal structure is not merely a reflection of descending motor commands or sensory (e.g., proprioceptive) feedback, but instead is an intrinsic property of the network. One of the key features of M1 population dynamics is that it exhibits low “tangling”^42^. This is taken as evidence for a first-order dynamical system, where the activity moves in a direction defined by the current activity state. Our study here strengthens those findings by challenging animals to move their neural activity in directions that would result in high tangling. The animals were not able to move their neural activity against a flow field despite strong incentives to do so. Instead, the neural activity continued to move in directions of low tangling.

In our previous work, we have examined which population activity patterns animals can readily produce^26,43,44^, and demonstrated how new population activity patterns that support new behaviors can form with extended training^27^. This study extends our previous finding by now addressing the temporal structure of neural population activity. The present work shows that neural activity is even further constrained than reported in earlier BCI studies: not only is neural activity constrained by an intrinsic manifold^26^ and neural repertoire^43^, we now show there are *temporal* constraints on neural population activity that are difficult to violate. It remains a possibility that with more time or practice, perhaps using incremental training^27^, the temporal structure of neural population activity might be altered. This process would presumably require altering the connectivity within M1 and/or how M1 is controlled by other parts of the brain. Another intriguing possibility is that even with extensive training, temporal structure would still persist. By this view, behavioral flexibility might be achieved by flexibly combining fairly fixed dynamical motifs^45,46^, rather than by forging new dynamics.

## Author contributions

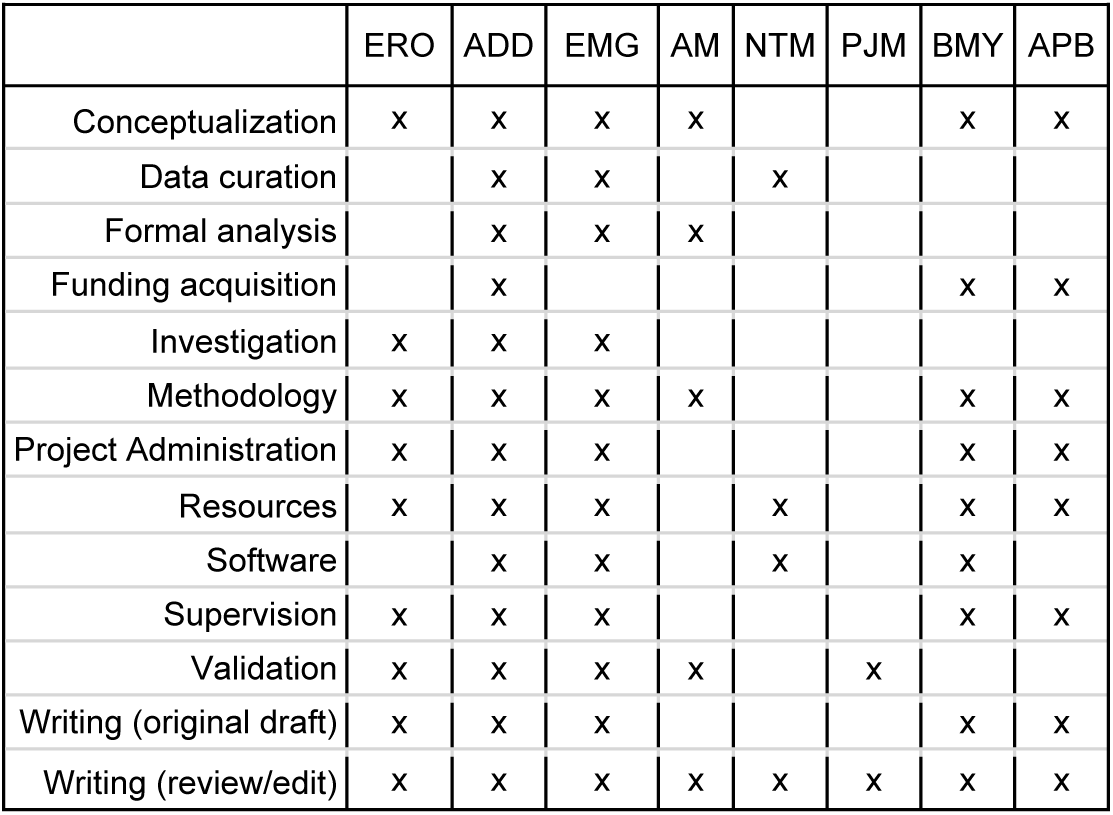

## Methods

All animal procedures were approved by the University of Pittsburgh Institutional Animal Care and Use Committee in accordance with the guidelines of the US Department of Agriculture, the International Association for the Assessment and Accreditation of Laboratory Animal Care, and the National Institutes of Health.

### Electrophysiology and behavioral monitoring

Prior to array implant surgeries, three animals were trained on a center-out reaching task. We implanted arrays in the motor cortex contralateral to the trained reaching arm. We recorded neural activity from the proximal arm regions of primary motor cortex (M1) in two male Rhesus macaques (monkey E, aged 12 years; monkey D, aged 12 years) using a 96-electrode (1 mm electrode length) microelectrode array (Blackrock Microsystems, Salt Lake City, UT, USA). In a third monkey, (monkey Q; male, aged 6 years) we implanted a 64-electrode (1.0 mm electrode length) array in the dorsal premotor cortex (PMd) and a 64-electrode (1.5 mm electrode length) array in M1. The neural signals from the electrodes were amplified and digitally processed using the TDT RZ2 system (Tucker Davis Technologies, Alachua, FL, USA). The digitized signals were bandpass filtered between 300 Hz and 3 kHz. We recorded neural units as threshold crossings, where a threshold crossing was detected when the depolarizing phase of the voltage signal crossed a threshold of 3 times the root-mean-square (RMS) voltage. We estimated the RMS voltage of the signal on each electrode prior to each experiment while the monkeys sat calmly in a darkened room. We recorded 94.0 ± 1.1, 79.9 ± 1.6, and 96.7 ± 0.5 neural units (mean ± standard deviation) from monkeys E, D, and Q, respectively. Neural activity was recorded 8-18 months after array implantation for monkey E, 9-18 months after implantation for monkey D, and 3-6 months after implantation for monkey Q.

During experiments, animals sat head-fixed (monkeys E and D) or head-free (monkey Q) in a primate chair in front of a visual display with both arms loosely restrained in a pronated posture. To reduce hand movements during the BCI trials and to ensure a consistent hand position, the monkeys placed their hand (contralateral to the array) on a horizontal “touch bar”. The touch bar was instrumented with either a contact sensor (monkey E) or a force transducer (monkeys E, D and Q, Mini40, ATI Industrial Automation, NC) to measure hand contact with the touch bar. Animals were required to maintain contact with the touch bar through the trial. Releasing the bar or exerting force outside of a predetermined window resulted in an immediate failure of the trial. The force window was calibrated relative to the weight of the hand on the force bar. Monkeys E and D had to maintain a force within a 11.8N window and monkey Q within a 1.3N window. The position of the hand (contralateral to the array) was tracked using an LED marker (Phasespace Inc., San Leandro, CA). There was minimal arm movement or force production during BCI trials (Extended Data Fig. 7)

### Behavioral tasks

For all behavioral tasks, trials were initiated by grasping and holding the touch bar for 250ms. Upon initiation, a BCI cursor and a target were displayed simultaneously on the monitor. The cursor remained fixed at the center of the workspace for 500ms (referred to as the “freeze period”), after which the cursor was placed under neural control. For all tasks, the animals were trained to move the cursor to acquire the presented target. The target was acquired when the cursor contacted the target (i.e., no hold time was enforced). Animals received a liquid reward upon successful completion of a trial. Each animal performed the following BCI tasks: center-out task, two-target task, grid task, intermediate target task and instructed path task. Each task is described in detail below.

#### Center-out task

During the center-out task, animals were required to move the BCI cursor from the center of the workspace to one of eight possible peripheral targets, arranged around a circle at 45 ° intervals. Peripheral targets were placed 90mm from the center of the workspace and were arranged in a radial pattern. Animals were given 4 seconds to acquire the peripheral target. Failure to acquire the target within this time or failing the touch bar conditions (i.e., releasing the bar or exerting force) resulted in a 2 second (monkeys D and Q) or a 5.5 second (monkey E) timeout. Targets were presented in a pseudo-random order such that each of the 8 targets was presented and attempted once before any target was repeated. A 160 trial block of the center-out task was used to calibrate the MoveInt decoder (see below).

#### Two-target task

The two-target task was a two-step task in which animals moved the BCI cursor to sequentially acquire two diametrically opposed peripheral targets (A and B). There are 4 possible target pairs: 0 & 180°, 90 & 270°, 45 & 225°, 135 & 315°. One target pair was tested in each session and was chosen before the start of the session. These peripheral targets were placed 90 mm from the center of the workspace. There were 9 experiments in which monkey Q was unable to acquire the peripheral targets at 90 mm, so we reduced the target distance to the greatest distance that the animal could acquire (80-85 mm). This task consisted of two steps. In the first step, the cursor and a peripheral target, (pseudo-randomly chosen to be target A or target B) simultaneously appeared on the screen and the monkey had 4 seconds to acquire the peripheral target (Extended Data Fig. 8A, left). We refer to the target acquired in step 1 as the “start target”. In the second step, the monkey had 4 seconds to move the cursor from the start target to the diametrically opposed target, i.e., from target A to target B or from target B to target A (Extended Data Fig. 8A, right). Successful acquisition of target B led to the animal receiving a liquid reward. Failing to acquire target B led to a 5.5 second penalty and all other failure modes resulted in a 2 second (monkeys D and Q) or a 3 second (monkey E) penalty.

#### Grid task

The grid task was a variant of the two-target task (see above) modified to include additional targets for the second step. We used the same peripheral target pair as was used for the two-target task in that session. As in the two-target task, the animal first acquired one of the two start targets, selected pseudorandomly. For the second step, there were three possible target locations: the diametrically opposed target or two targets perpendicular to the target pair (Supp Fig. 8B). The probabilities of the targets were weighted so that there were 100 total trials to the diametrically opposite target and a 40 total trials to the other two targets. For each step, the animal had 4 seconds to acquire the target and no target hold time was enforced. Following successful completion of the second step, the animal received a liquid reward.

#### Intermediate target task

The intermediate target task was a two-step target acquisition task, where the first step was the same as in the two-target task (see above). The second step was to acquire was an “intermediate target” (IT), placed along an axis orthogonal to the target pair axis (Fig. 6). The location of the intermediate target was selected for each experiment so that the animal could acquire it from both target A and target B (Extended Data Fig. 8C). We positioned the IT as close as possible to the path defined by the observed cursor trajectory for one direction (e.g., A-to-B), while still being able to acquire from the opposite target condition (e.g., B-to-A). To determine the intermediate target position, we gradually increased the target distance from the center of the workspace in 10% increments of the peripheral target distance until the success rate began to decline. The final IT position was chosen to ensure both of the following: 1) a high success rate from both start targets was maintained and 2) the target location was aligned with the path of the flow field (blue arrow in Extended Data Fig. 8C). Across experiments, this procedure resulted in an IT position that was 31.2 ± 11.4 mm from the center of the workspace. The animal performed approximately 100 total trials of the intermediate target task, moving from either start target to the IT (i.e., A-to-IT, B-to-IT). These trials were used to estimate trajectory flexibility (see ‘Estimating trajectory flexibility’, below).

#### Instructed path task

The instructed path task was a variant of the intermediate target task (see above), where the cursor path was instructed with a visual boundary around the start target and the intermediate target (Fig. 7A). In order to succeed at the task, the animals were required to keep the cursor within the boundary as they acquired the intermediate target. The boundaries were straight capped cylinders that encased the allowable path between the start target and the intermediate target. The boundary edges were equidistant (ranging from 30-130mm) from the target axis (i.e., the line that connects the targets) and rounded near the two target positions.

An instructed path trial began with the presentation of the start target. After successful acquisition of the start target, the intermediate target and the visual boundary appeared simultaneously. Animals then had 4 seconds to acquire the intermediate target (i.e., the “acquire time”) without touching the visual boundary to receive a reward. Failing the boundary requirement resulted in a time penalty that was 2 seconds plus the remaining acquire time. This trial structure ensured that animals would receive shorter time penalties for trials in which they were actively attempting to acquire the target and longer penalties for failing quickly.

The instructed path task started with a boundary size that required minimal modification of cursor trajectories for success. The first boundary was chosen to have a width of 110 or 120 mm, with the exception of a single session for monkey D (150mm width) and two sessions for monkey E (80mm width). Then, we gradually reduced the size of the boundary to encourage the animals to modify their cursor trajectories. The reduction happened in one of two ways. For all sessions with monkeys E and D, and 6 of 13 sessions with monkey Q, we evaluated the animal’s task performance every 25 trials and reduced the boundary’s width by 10 mm if the animal exceeded a 75% success rate. For the other 7 of 13 sessions of monkey Q, we evaluated the animal’s task performance every 25 trials and reduced the boundary size such that new smaller size would yield a predicted 75% success rate. With both approaches, if the actual success rate failed to meet the 75% success rate threshold, we began evaluating performance in 50 trial blocks. If the success rate threshold was not met in a given block, the boundary size would stay the same for another 50-trial block. The boundary was not increased once reduced, except in rare instances where the initial boundary width was too difficult for the animal. The animal performed a minimum of 500 instructed path trials in experimental sessions that included this task (Extended Data Fig. 5). For the final 100 instructed path trials, we kept the task parameters constant even if the animal met the success rate threshold. Experimental sessions which included the instructed path task are summarized in Supp. Table 1.

### BCI mappings

To study temporal constraints on neural population activity, we sought to give animals moment by moment feedback of their neural trajectories. To accomplish this, the recorded neural activity was transformed to the position of a computer cursor. We used Gaussian-process factor analysis (GPFA) to transform the ∼90D recorded neural activity at each time step into a *p*-dimensional latent state^28^. For all experiments, we used *p* = 10 dimensions, as this has been found to capture most of the shared variability in motor cortex during BCI control^26^.

Ideally, we would provide the animal visual feedback of all 10 dimensions, but providing 10 dimensions of feedback is challenging to set up experimentally. Instead, we provided visual feedback as a particular 2D projection of the 10D latent state. The 10D latent states were mapped to 2D cursor position via a BCI mapping. We used two types of BCI mappings defined below: movement intention (MoveInt) and separation maximizing (SepMax). Unlike our previous BCI studies^26,27^, which mapped neural activity to cursor *velocity*, here neural activity is mapped to cursor *position* to establish a direct correspondence between the neural activity patterns and workspace location.

### Extracting neural trajectories

To calculate spike counts, threshold crossing events for each neural unit were binned in non-overlapping 45ms time windows. We used GPFA to extract the latent state at a given point in time ***z***_*t*_ ∈ *R*^10×1^ from the spike counts ***u***_*t*_ ∈ *R*^*q*×1^ at the recent past and the current time points, where *q* is the number of neural units. GPFA defines a linear-Gaussian relationship between latent states and spike counts as

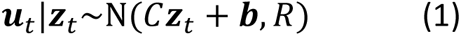

where *C* ∈ *R*^*q*×10^ specifies the relationship between the latent states and spike counts, ***b*** ∈ *R*^*q*×1^ is the mean activity of each neural unit, and *R* ∈ *R*^*q*×*q*^ is a diagonal matrix specifying the independent variance of the neural units.

Latent states are related across time using Gaussian processes. The neural trajectory for the *i*^th^ latent state at time steps 1 to *T*, ***z***_*i*_ = [*Z*_*i*,1_, …, *Z*_*i*,*T*_], is defined as

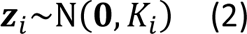

where *K*_*i*_ ∈ *R*^*T*×*T*^ is a covariance matrix defining the relationship between the *i*^th^ latent state at different points in time and *i* = 1, …, 10.

The GPFA parameters *C*, ***b***, *R*, and latent timescales in *K*_*i*_ were fit using the expectation-maximization algorithm as described in Yu et al., 2009^28^. For monkey E, a GPFA model was fit to the center-out trials to define the MoveInt mapping and a separate GPFA model was fit to the two-target task trials to define the SepMax mapping (see Movement Intention Mapping and Separation-Maximizing Mapping, below). For monkeys D and Q, a single GPFA model fit to the center-out trials was used to define the MoveInt and SepMax mappings. We included neural activity from target onset to target acquisition on successful trials and neural activity from target onset to moment of failure on failed trials.

To extract the neural trajectories, first we extracted the unsmoothed latent state,

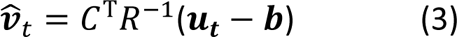

where ***v̂***_*t*_ ∈ *R*^10×1^. The standard form of GPFA uses both past, current, and future neural activity to estimate the current neural state^28^. Since we are presenting neural trajectories in real time to the animals, we are limited to using past and current neural activity. We therefore designed a causal implementation of GPFA^47^ in which the latent state at the *t*^th^ time step is only determined by neural activity from the previous seven time steps (i.e., *t* − 6 to *t*, approximately 315ms into the past), rather than neural activity from all past and future times. We concatenated the unsmoothed latent states for the previous 7 time steps

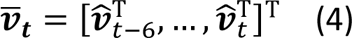

Where ***v̄***_***t***_∈*R*(7∗10)×1 Finally, the neural trajectories were extracted in real-time as

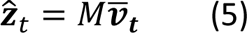

where ***Ẑ**_t_*∈*R*10×1 is the estimate of the latent state at time *t* and *M* ∈ *R*^10×^(7∗10) is a ‘smoothing matrix’ describing the contribution of past and current spiking activity on the latent state at time *t* (Extended Data Fig. 9). The contributions corresponding to time steps *t* − 7 and beyond were negligible relative to the contributions from more recent time steps (Extended Data Fig. 9). The smoothing matrix *M* is obtained from the model parameters *C*, *R*, and *K*_*i*_ (*i* = 1, …, 10) as described in Yu et al., 2009^28^. The spike count history used to extract neural trajectories was reset to zero at the beginning of each trial. There were no “edge effects” of estimating ***ẑ***_*t*_ related to this reset, as the touch bar hold time (250ms) and the cursor freeze period (500ms) occurring at the beginning of each trial were longer than the 315ms spiking history used for the causal GPFA mapping.

Using the ***ẑ***_*t*_ extracted by GPFA, we specified a BCI mapping of the form

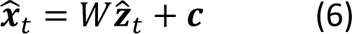

to convert ***ẑ***_*t*_ into a cursor position ***x̂***_*t*_∈*R*^2×1^, where *W* ∈ *R*^2×10^ is a weight matrix, and ***c*** ∈ *R*^2×1^ is a positional offset. Each BCI mapping (MoveInt and SepMax) used a different *W* and ***c***, defined below.

### Movement intention mapping

Each experiment began with calibrating the movement intention (MoveInt) mapping. We used a gradual training process^26^ to determine the parameters of the this mapping. This process consisted of a series of five 32-trial blocks of center-out trials, in which we updated the mapping parameters after each block, using all accumulated data up to that point. The first block consisted of passive observation trials, during which the cursor was moved to 8 peripheral targets under computer control. The cursor was moved at constant velocity (0.15 m/s) straight to the target. The targets were presented in a pseudo-random order (∼4 trials per target). Following the observation block, animals were given control of the computer cursor, but we attenuated the perpendicular error in order to discourage online movement corrections. We reduced the amount of perpendicular error attenuation each block so that the animal had full online control of the cursor by the final training block. After the final block, we used the data from all five blocks to calibrate the MoveInt mapping.

The MoveInt BCI mapping was designed to provide animals with proficient cursor control, such that they were able to move the cursor quickly and accurately to targets placed throughout the workspace. To identify the MoveInt mapping we used linear regression (Equation 6) to solve for the parameters *W*_*MI*_ ∈ *R*^2×10^ and ***c***_*MI*_ ∈ *R*^2×1^ that best predicted the assumed intent of the animals given their neural trajectories. More specifically, mapping parameters were defined as

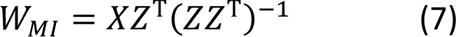

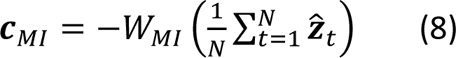

where *Z*=[***ẑ***_1_,…,***ẑ***_*N*_]∈*R*^10×*N*^ comprises the latent states estimated by GPFA and *X* = [***X***_1_, …, ***X***_*N*_] ∈ *R*^2×*N*^ comprises the associated intended cursor positions (see below) and *N* is the total number of time steps during the calibration trials.

The calibration data for each trial consisted of sets of {***ẑ***_*t*_, ***X***_*t*_} for two time epochs: the first epoch comprised 3 time bins of activity within each trial that captured baseline activity ( *T*_*base*_) and the second epoch comprised 5 time bins of activity within each trial that captured activity when the animal was engaged in the task (*T*_*engaged*_). The details of which time bins comprised each epoch varied for each monkey and are described below. For all animals, *N* = *n* ∗ (*T*_*base*_ +*T*_*engaged*_), where *n* is the number of calibration trials. Therefore, after each 32-trial block, *N* increased by 256 (i.e., 32*(3+5)) time steps.

For monkey E, the first time epoch consisted of the first 3 time bins (approx. 135ms) of the freeze period. During this period, we assumed that the animal was not attempting to move the cursor and set ***X***_*t*_ = [0 *mm*, 0 *mm*]^T^. The second time epoch was 5 time bins in duration (approx. 225ms) immediately preceding acquisition of the BCI target. During these time bins we assumed that the animal was intending that the cursor be placed at the target location (e.g., ***X***_*t*_ = [90 *mm*, 0 *mm*]^T^ for rightward targets). We found that these assumptions generally worked well for monkey E, whose neural activity was highly stereotyped during calibration.

For monkey D, we found that using the above calibration approach led to MoveInt mappings that did not provide good cursor control (i.e., the animal was not able to consistently acquire all targets). This is likely due to neural responses in the 5 time bins preceding acquisition of the target being more variable on a trial-to-trial basis than we observed for the other monkeys.

Instead, we identified target intent for monkey D using neural activity during the freeze period, rather than activity immediately preceding the acquisition of the BCI targets.

Monkey Q adjusted his grip on the force bar between each trial which led to some inconsistencies at the beginning of the trial until he settled into a stable grip for the remainder of the trial. To account for this behavioral variability at the beginning of the trial, we defined the first epoch as the last three time bins of the freeze period, rather than the first three time bins of the freeze period. The second epoch was defined in the same way as for monkey E.

### Separation-maximizing mapping

The separation-maximizing (SepMax) BCI mapping was designed to highlight projections of neural activity in which neural trajectories took different paths through the latent space when moving between target pairs in the two-target task. We identified projections that satisfied three objectives (Extended Data Fig. 10): 1. Maximization of the separation between the midpoints of the A-to-B trajectories (***z̄***_*AB*_) and the B-to-A trajectories (***z̄***_*BA*_), 2. Minimization of the trial-to-trial variance at the midpoints (∑_*AB*_ and ∑_*BA*_), and 3. Maximization of the distance between the starting points of the A-to-B trajectory (***z̄***_*A*_) and the B-to-A trajectory (***Z̄**_B_*).

We first computed the trial-averaged starting points ***z̄***_*A*_ and ***z̄***_*B*_ ∈ *R*^10×1^ for the A-to-B and B-to-A trajectories, respectively (Extended Data Fig. 10A). We then defined an axis connecting ***z̄***_*A*_ and ***Z̄**_B_*, along with its midpoint

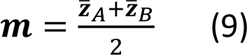

where ***m*** ∈ *R*^10×1^. For a given single-trial neural trajectory, we projected each of its latent states onto this axis (Eqn. 5). We defined the midpoint of this neural trajectory as ***ẑ***_*tc*_, where *t*_*c*_ is the time point at which the projection of ***ẑ***_*t*_ onto the axis was closest to ***m***. The trial averages of ***ẑ***_*t*_ for the A-to-B trajectories and the B-to-A trajectories are ***z̄***_*AB*_ ∈ *R*^10×1^ and ***z̄***_*BA*_ ∈ *R*^10×1^, respectively (Extended Data Fig. 10B). The covariance across trials of ***ẑ***_*t*_ for the A-to-B trajectories and the B-to-A trajectories are ∑_*AB*_ ∈ *R*^10×10^ and ∑_*BA*_ ∈ *R*^10×10^, respectively.

To identify the SepMax projection, we used an optimization procedure^48^ to identify a set of basis vectors to project the 10D neural trajectories into 2D cursor positions to satisfy the objectives above. Specifically, we sought to find an orthonormal set of vectors *P*_*SM*_ = [*p*_1_, *p*_2_] ∈ *R*^10×2^ which minimized the objective function:

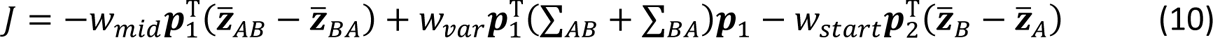

where *w*_*mid*_, *w*_*var*_, and *w*_*start*_ are scalar weighting factors. The first term of the objective function maximizes the midpoint separation, the second term minimizes the trial-to-trial variance, and the third term maximizes the starting point separation. Thus, *p*_1_ is the dimension along which the A-to-B and B-to-A trajectories are separated, and *p*_2_ is dimension along which the targets are separated (i.e., the target axis).

Weighting factors *w*_*mid*_, *w*_*var*_, and *w*_*start*_ were used to specify the relative influence of the midpoint, covariance, and starting point terms on the overall objective function value. For monkey E, each of these terms was set to 1. For monkeys D and Q, the weighting factors were chosen such that the identified projections were not dominated by any single objective. To find the weighting factors each term in the objective function (ignoring *w*_*mid*_, *w*_*var*_, and *w*_*start*_) was calculated for 10,000 different random orthonomal projections *P*_*SM*_. We set each weighting factor to be the inverse of the range (i.e., maximum value – minimum value) of that term.

Objective function minimization proceeded as outlined in Cunningham and Gharamani, 2015^48^, with a convergence criteria of Δ*J* = 10^−10^, a maximum of 1000 gradient iterations and a line search step size of 0.1. The SepMax mapping was calculated from the 160 trials of the two-target task, or from the ∼100 trials of the grid task to the diametrically opposed targets.

To make BCI control with the separation-maximizing projection as intuitive as possible for the animal, we made the visual feedback as consistent as possible between the different BCI feedback conditions. To determine the SepMax mapping parameters (*W*_*SM*_ and ***c***_*SM*_), we aligned the space defined by *P*_*SM*_ with the animals’ workspace such that the starting points of the A-to-B and B-to-A trajectories in the SepMax mapping were at the same position as targets A and B in the cursor workspace. Specifically, we defined

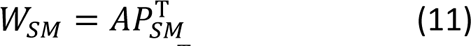

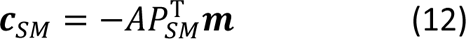

where *W*_*SM*_ ∈ *R*^2×10^, *A* ∈ *R*^2×2^, ***c***_*SM*_ ∈ *R*^2×1^, and ***m*** is defined in Eqn. 9. The matrix

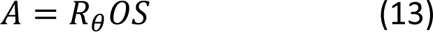

specifies a linear transformation involving three operations to provide intuitive visual feedback with the SepMax mapping. First, we scale the axes of *P*_*SM*_. The scaling matrix *S* = *sI* ∈ *R*^2×2^ is

a diagonal matrix that scales the axes of *P*_*SM*_, such that the distance between ***z̄***_*A*_ and ***z̄***_*B*_ is equal to the distance between targets A and B in the MoveInt mapping. Then, we optionally flip the projection about the target axis using the matrix *O* ∈ *R*^2×2^ (Extended Data Fig. 10C, D). We want to orient the SepMax projection such that attempted movements move the cursor in the expected direction (e.g., when the monkey intends to move up, the cursor moves up rather than down). To determine the sign of *p*_1_that achieves this goal, we visually inspected the neural trajectories during the grid task in both orientations (Extended Data Fig. 8B, C). We chose the sign of *p*_1_ such that the endpoints of the neural trajectories in the SepMax projection were closest to the associated target location (Extended Data Fig. 10C). This choice reduced the cognitive burden imposed on animals when using the SepMax mapping. Finally, we rotated *p*_2_so that it aligned with the workspace targets A and B. The matrix *R*_*θ*_ ∈ *R*^2×2^ rotates the projection through angle *θ*, where *θ* is the angular difference between the axis connecting the workspace targets in the MoveInt mapping and the *p*_2_ axis.

### Experimental flow

The experimental flow for a single session was the same for all monkeys. Each experiment began by calibrating a MoveInt mapping that captured the animal’s movement intention during the center-out task. The MoveInt mapping was then used during the two-target task or the grid task to identify the SepMax mapping for one target pair (Fig. 3). Then, we tested the flexibility of the temporal structure captured by the SepMax mapping with three experimental manipulations. First, we gave the animal visual feedback of the dimensions where temporal structure was evident by having the animal perform the two-target task using the SepMax mapping (Fig. 4). Then, we used the intermediate target task to ask if the animal could produce time-reversed neural trajectories (Fig. 6). Finally, in the instructed path task we directly challenged the animal to follow a prescribed path (Fig. 7).

The details of how we identified the SepMax projection varied somewhat for each monkey. For the majority of sessions for monkeys E and D, only a single target pair was tested. We selected the target pair to be tested pseudo-randomly before the start of the experiment. We identified the SepMax projection for that target pair using either 160 trials of the two-target task or 140 trials of the grid task with the MoveInt mapping. If we used the grid task, only the ∼100 trials to the selected target pair were used to identify the SepMax projection. For monkey Q, we used a different procedure because he showed high variability of neural trajectory separation across sessions. We selected the target pair to be tested each day from 160 two-target trials comprising all four target pairs (i.e., 40 trials per target pair) using the MoveInt mapping. The selection criterion balanced the desire to test each target pair during multiple sessions while also prioritizing target pairs with strong trajectory separation. After selecting the target pair to be tested, we identified the SepMax projection using the ∼100 trials of the grid task to the selected target pair.

Once we identified the SepMax projection, the animals exclusively used the SepMax mapping to control the cursor for the remainder of the behavioral tasks in the session. To assess the persistence of temporal structure when it was provided as visual feedback to the monkey, the animal performed 100 trials of the two-target task using the SepMax mapping (Fig. 4).

Next, we ran 50-100 trials of the intermediate target task for each start target using the selected target pair while we adjusted the intermediate target position. We used these trials to establish the location of the intermediate target. After setting the position of the intermediate target, we ran an additional 100 trials of the intermediate target task with both start target positions. These trials were used to estimate trajectory flexibility (Fig. 6, see ‘Estimating trajectory flexibility’, below). For the final task of each session, the animal performed 500 trials of the instructed path task. We reduced the visual boundary according to animal’s success rate, as described above (Fig. 7, see ‘Instructed path task’).

We analyzed 111 two-target sessions with the SepMax mapping (50, 40, and 21 sessions for monkeys E, D, and Q, respectively). A subset of those sessions also included the intermediate target and the instructed path tasks (28, 9, and 13 sessions for monkeys E, D, and Q, respectively). We analyzed data from 18 additional sessions in which the SepMax projection was reflected (Extended Data Fig. 4). We excluded a session if any trials were corrupted or lost during the data saving process, or if animal motivation issues prevented us from obtaining a MoveInt mapping that provided satisfactory control. Overall, this exclusion process resulted in excluding 2 sessions due to lost or corrupted date and 4 sessions due to low motivation.

### Analyses Discriminability index

We sought to measure how distinct the neural trajectories were for different conditions. We used a discriminability index (*d*′) to measure the separation of the midpoints of the A-to-B versus B-to-A neural trajectories (Fig. 3; see Separation-maximizing mapping). We defined a unit vector pointing between the midpoint of the A-to-B and B-to-A trajectories

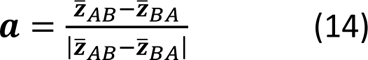

where ***a*** ∈ *R*^10×1^ (Extended Data Fig. 10E). For each trial, we projected the midpoint of the neural trajectory ***ẑ***_*tc*_ onto ***a***. Then we determined the mean and variance across trials of these projections, separately for the A-to-B and B-to-A trajectories. These means and variances were used to calculate *d*′

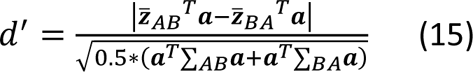

Larger values of *d*′ correspond to latent states that are more separable between the A-to-B and B-to-A conditions.

### Flow field analysis

We used a flow field analysis to compare neural trajectories in different 2D projections across experimental conditions. The flow field estimates the velocity as a function of position. In other words, this technique sought to estimate ***x***_*t*+1_ − ***x***_*t*_ = *f*(***x***_*t*_) where ***x***_*t*_ ∈ *R*^2×1^ is a 2D projection of ***ẑ***_*t*_ from Eqn. 5. In Fig. 4, we characterize the flow fields of the cursor trajectories, i.e., the neural trajectories in the MoveInt and SepMax projections. In Extended Data Fig. 3, we characterize the flow fields of the neural trajectories in random 2D projections.

To estimate the flow field in a given 2D projection, we first partitioned the 2D space into a set of square voxels (Extended Data Fig. 3A). We then calculated the velocity (i.e., ***x***_*t*+1_ − ***x***_*t*_) of the neural trajectory in the 2D space at each time point on individual trials. For each voxel, we averaged the velocities of the latent states that were located within that voxel (Extended Data Fig. 3B, C). We used a voxel size of 20 mm for the MoveInt and SepMax projections. Average velocity vectors for a given voxel were only considered valid if there were a minimum of 2 time points that were located within that voxel. We calculated a separate flow field for each target condition (i.e., A-to-B and B-to-A) to capture how the neural trajectory unfolds from a given initial condition. The flow fields for each condition are plotted together to visualize the overall flow (Fig. 5A). The separate flow fields for each target condition were averaged to create a single flow field for a given projection. To compare two flow fields (Fig. 5B and Extended Data Fig. 3D-H), we calculated the mean squared difference between velocity vectors of corresponding voxels of the flow fields. Then we took the median of those values across voxels.

### Estimating trajectory flexibility: initial angle

To assess trajectory flexibility, we first measured the initial angle for cursor trajectories during the two-target task. This angle reflects how the trajectories emanate from the start targets as captured by the flow field in Fig. 5. We sought to assess the extent to which the heading direction of cursor trajectories could be altered by the animal in the intermediate target (Fig. 6) and instructed path tasks (Fig.7). We compared the initial angle of the cursor trajectories during the intermediate target and the instructed path tasks to the initial angle during the two-target trials. If the initial angles are the same, it would suggest that the activity time courses are not flexible. However, if the initial angles for the intermediate target or the instructed path tasks were smaller than those for the two-target task, it would indicate that the activity time courses are flexible and that the flow field can be violated.

In the intermediate target and instructed path tasks, we measured the unsigned “initial angle” between the heading direction of the trajectory and the vector pointing from the start target to the intermediate target (i.e., the direct path). We defined the heading direction of each trajectory as the vector from the first to the fourth time point of the cursor trajectory. The fourth time point (180 ms) was chosen because it is late enough to ensure that the animal was responding to the visual display of the intermediate target, but early enough to minimize the effect of any error corrections that occurred later in the trial. The initial angle was computed for each trial and then averaged across successful trials and failed trials that reached at least the fourth time point. Including failed trials helped to characterize dynamical constraints for the instructed path task, in which reducing the size of the boundary led to lower success rates, without over-representing successful trajectories that might have been more direct to the intermediate target.

To understand to what extent the cursor trajectories during the intermediate target and instructed path tasks were consistent with the flow field defined by the two-target trajectories, we also computed the initial angle of the two-target trials as a reference, using the same method as described above. We defined the initial angle of the two-target trials as the average initial angle of the first 20 trials from the same start target as was tested in the intermediate target and instructed path tasks i.e., the early two-target trials. Using the early two-target trials allowed us to construct control comparisons (described in detail below) with the same reference.

To compare the change in initial angle across experiments, we normalized the change in initial angle:

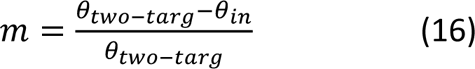

where *θ*_*two*−*targ*_ is the trial-averaged initial angle for early two-target trials, and *θ*_*in*_ is the trial-averaged initial angle for the intermediate trials (Fig. 6) or instructed path trials (Fig. 7) defined relative to the direct path to the intermediate target. A value of *m* = 0 means that there is no change in initial angle relative to the two-target trials. A value of *m* = 1 means the animal is able to move the cursor straight from the start target to the intermediate target along the direct path, i.e., *θ*_*in*_ = 0.

For reference, we compared *m* to a “no change” condition and a “full change” condition. We constructed the no-change condition using trials in which there is no expectation that the initial angle of the trajectories should change (Fig. 6F). We compared the change in initial angle between the first 20 (i.e., early) trials and the last 20 (i.e., late) trials from the same start target of the two-target task with the SepMax mapping. Using the early two-target trials allowed us to make all the initial angle comparisons to the same reference.

We constructed the full-change condition using trials in which the animal demonstrated flexible control. To do so, we used grid task trials because in the grid task the animal could move directly to different instructed targets from the same start target using the MoveInt mapping (Fig. 6H). We computed the initial angle for the A-to-B trajectories, as well as the initial angle for the trajectories from target A to the perpendicular grid target that was in the same direction as the intermediate target. Then we computed the change in initial angle

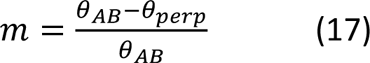

where *θ*_*AB*_ is the trial-averaged initial angle for A-to-B trials, and *θ*_*perp*_ is the trial-averaged initial angle for the target A to perpendicular grid target trials. Note that this full change condition reflects not only the flexible control that the animal has with the MoveInt mapping but also some behavioral idiosyncrasies. Animals did not always change their initial angle to move directly to the perpendicular grid targets, rather they would sometimes follow the path of the A-to-B trajectories before correcting to the perpendicular target. This resulted in smaller changes in the initial angle metric.

---

**Extended Data Figure 1.**
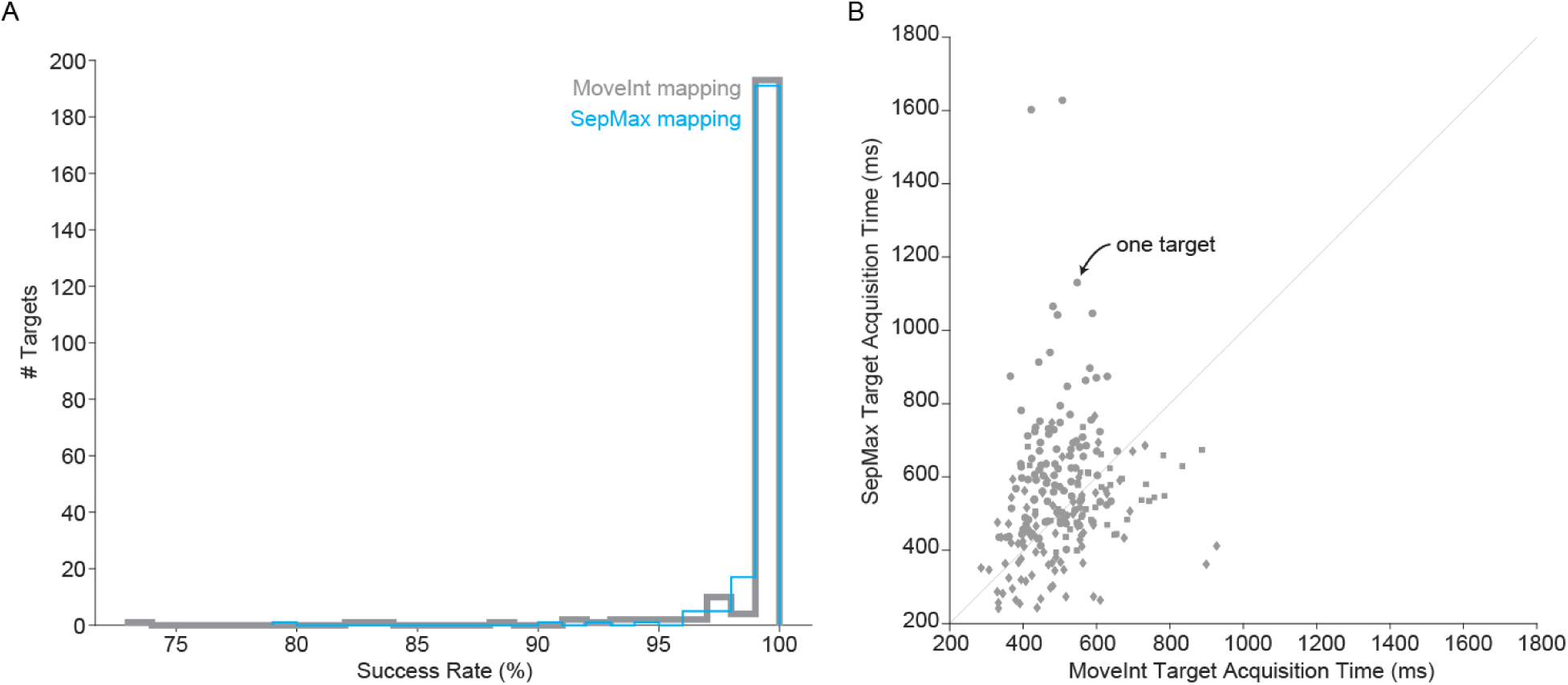
BCI performance is similar regardless of the projection providing feedback. BCI performance with the separation-maximizing mapping is similar to the performance with the movement intention mapping. We quantified performance in the two-target task using (A) success rate and (B) target acquisition times. N = 222 targets (111 sessions, 2 targets per session). A. The animals are highly proficient at the two-target task with both mappings, nearly always performing at 100%. Success rate is calculated for A-to-B and B-to-A movements separately. B. Average target acquisition times as the monkey used the MoveInt mapping (horizontal axis) and the SepMax mapping (vertical axis). The target acquisition time is the time it takes the monkey to move the cursor in step 2 of the two-target task (see Methods) and is calculated separately for A-to-B and B-to-A movements. Each dot represents one target. Across all monkeys, the acquisition times with the MoveInt mapping were 507.8 ± 109.5 ms (mean ± standard deviation) and those with the SepMax mapping were 553.4 ± 188.1 ms.

**Extended Data Figure 2.**
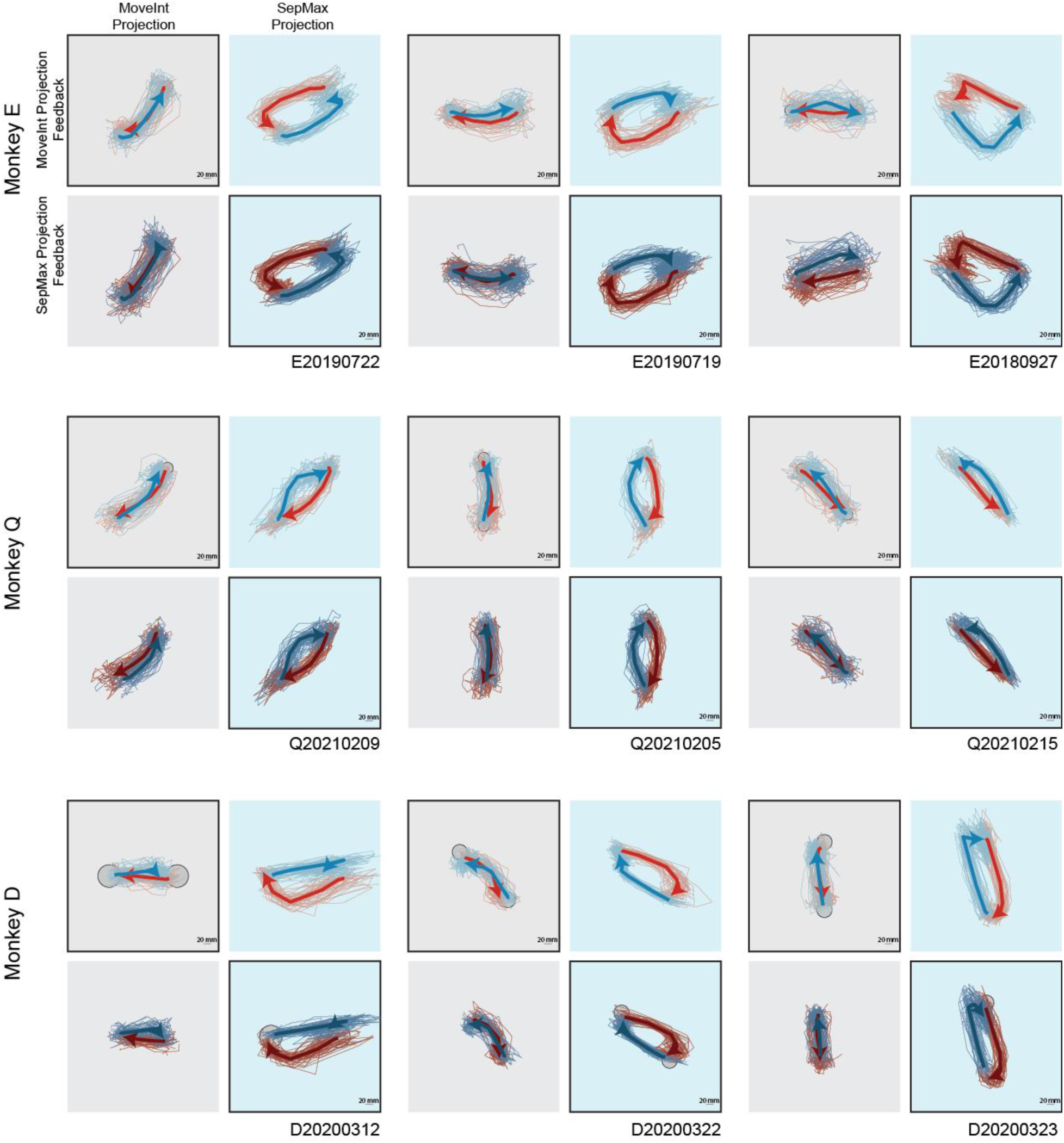
The persistence of temporal structure in neural population activity is robust. Here we show representative example sessions from each monkey and different target pairs. Trajectories are plotted in the MoveInt (gray background) and SepMax (light blue background) projections. When a projection is providing feedback to the monkey for BCI control, the subpanel has a black outline. When a projection is unseen by the monkey, the subpanel does not have an outline. Same conventions as Fig 4D.

**Extended Data Figure 3.**
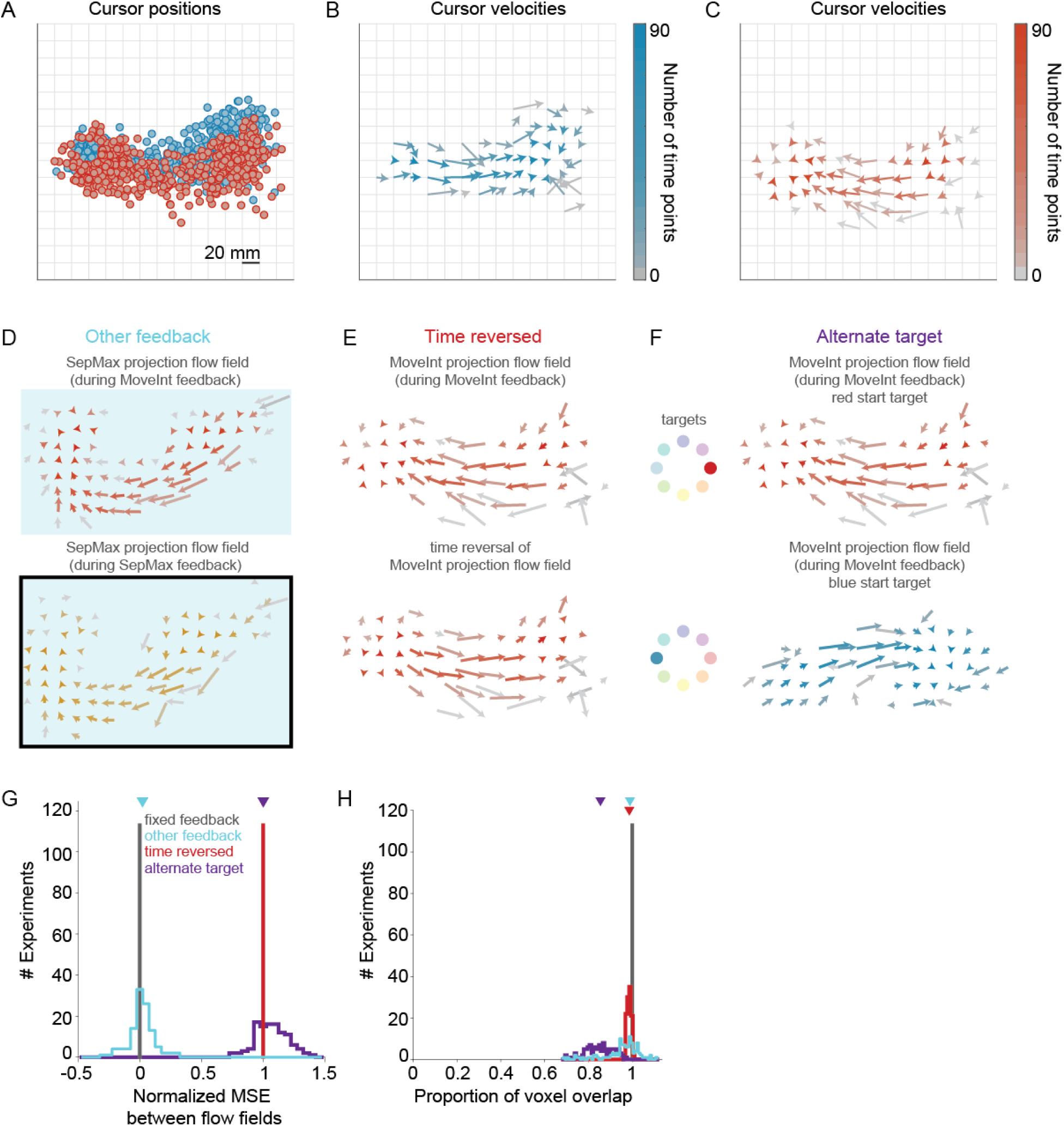
Flow fields. We used a flow field analysis to compare neural trajectories in different 2D projections. A. To determine a cursor trajectory flow field, we segmented the 2D workspace projection into a grid of 20 mm voxels. Dots indicate cursor positions at each time point for all trials. Dots are colored by the start target (blue: start target A at left of workspace; red: start target B at right of workspace). B, C. The velocity for a given voxel is defined as the velocity (***x̂***_*t*+1_ - ***x̂***_*t*_) averaged across all time points with a cursor position (***x̂***_*t*_) in that voxel. For visual clarity, we show the flow fields separately for each target condition (panel B: target A to target B, panel C: target B to target A). The length of the arrows represents the magnitude of the velocity and the orientation of the arrows indicates the direction. The color indicates the number of time points that contributed to the average. D-H. The flow field analysis (Fig. 5) shows that the time courses of neural activity are strongly constrained within the SepMax projection, regardless of whether the animal receives feedback of their neural activity in the MoveInt or SepMax projections. However, it is not yet clear whether these constraints are limited to specific subspaces, or whether neural trajectories are constrained in all dimensions of the 10D space. To test this, we applied a flow field analysis similar to that used in Fig. 5 (see Methods and panels A-C above) to neural activity in random 2D projections of the 10D space. We first projected neural trajectories into random 2D subspaces:

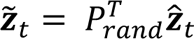 where *P*_*rand*_ ∈ *R*^10×2^ is a random matrix with orthonormal columns and ***ẑ***_*t*_ is the latent state at time step *t* as defined in Eqn. 5. Then we estimated flow fields in this 2D subspace, using a 1×1 latent unit voxel size. D. “Other feedback” comparison. We compared the flow fields in a given projection between feedback conditions. For example, in the SepMax projection we compared the flow field during MoveInt feedback (top) and to the flow field during SepMax feedback (bottom). Note that the illustrated flow field comparison is the same as is shown in Fig. 5 for the SepMax projection (Fig. 5 light blue arrow), but we repeat the comparison for 400 random 2D projections per experiment to get the cyan distribution in (G) and (H). In order to appreciate the amount of change we observe in the flow fields in the “Other feedback” comparison, we constructed control distributions for which we expect no change and maximal change in the flow fields. For a no-change distribution, we compared flow fields for different subsets of trials with the same visual feedback. We call this distribution “Fixed feedback.” For maximal change distributions, we constructed two distributions: one in which the flow fields are overlapping and maximally different, i.e., the “Time-reversed” condition (E), and one in which the flow fields are different but less overlapping, i.e., the “Alternate target” condition (F). E. “Time-reversed” comparison. In the Time-reversed comparison, we compared the flow fields between trials for a given feedback condition, e.g., MoveInt trajectories (top) to a time-reversed version of the MoveInt trajectories (bottom). We generated the time-reversed neural trajectories in an offline analysis by reversing the temporal sequence of trajectories ***ẑ***_*t*_, making the last time point the first and the first time point the last. Note that the schematic simply reverses the direction of the velocity vectors, rather than illustrating the time-reversed flow field. F. “Alternate target” comparison. In the Alternate-target comparison, we compared the A-to-B flow field to the B-to-A flow field for a given feedback condition. For example, we compared trajectories from one start target (top) to trajectories from the other start target (bottom) during MoveInt feedback. G. Quantification of flow field comparisons. We compared 400 random 2D projections per experiment for each flow field comparison. By comparing the difference in flow fields for the “Other feedback” comparison to these three control distributions across random projections, we can determine whether the feedback provided to the monkeys changed neural trajectories in the full 10D space. For each session, we compare the flow fields of 50 random trial splits in each of 400 random projections. The total number of available trials for a given condition (49 ± 3.8 trials) was randomly sub-selected to form two sets of 20 trials and flow fields were estimated for each set. All comparisons were between flow fields for a given random split of trials (with the exception of the fixed feedback case). We calculated the mean squared error (MSE) between velocity vectors of corresponding voxels of the flow fields and took the median of those values across voxels (see Methods) for each of the random trial splits in each projection. We quantified the flow difference, *m*_*j*_, as the mean of the distribution of MSE across trial splits for each projection. To compare these distributions across sessions, we normalized the flow difference with respect to the Fixed-feedback as the lower limit, and Time-reversed as the upper limit

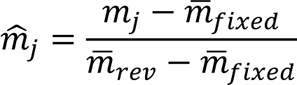 where *m̂*_*j*_ is the normalized flow field difference for *j*th projection, *m*_*j*_ is the flow difference for the *j*th projection, *m̄*_*fixed*_ and *m̄*_*rev*_ are the per-session average across projections of the flow difference magnitude for the Fixed-feedback and Time-reversed distributions, respectively. A *m̂*_*j*_ = 0 indicated that there was no change in flow difference magnitude between distributions, while *m̂*_*j*_ = 1 indicated that flow difference magnitudes were maximally different between comparison conditions. We averaged *m̂*_*j*_ across projections to yield a single value, *m̂*, for each session. By definition, *m̂* = 0 for the Fixed feedback comparisons and *m̂* = 1 for the Time-reversed comparisons. We found that the flow difference for the Other feedback comparisons were small. The Other feedback (cyan) comparison was not significantly different from the Fixed feedback (gray) comparison (paired t-test, p=0.0934). H. We also measured “flow field overlap,” which quantifies the degree to which the trajectories occupy the same region of state space. Flow field overlap, *o*_*i*_ was quantified as the number of voxels with a minimum of 2 time points within that voxel for each of the flow fields being compared. Like the flow difference metric, we calculated flow field overlap of 50 random trial splits for each of the 400 random projections. To compare these distributions across sessions, we normalized the flow field overlap with respect to the Fixed feedback comparison which has the highest degree of observed flow field overlap

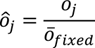 where *ô*_*j*_ is the normalized flow field overlap for *j*th projection, *o*_*j*_ is the flow field overlap for the *j*th projection, and *o*_*fixed*_ is the per session average across projections of the overlapping voxels for the fixed feedback distributions. A *ô*_*j*_ = 0 indicates that the region of the state space occupied by the trajectories was highly non-overlapping between distributions, while *ô*_*j*_ = 1 indicates that the overlap between trajectories was the same as the amount of overlap observed in the fixed feedback condition. We averaged *ô*_*j*_ across projections to yield a single value, *ô*,for each session. We found that the Fixed feedback, Other feedback and Time-reversed comparisons all show high flow field overlap, although the flow field overlap for the Fixed feedback comparison was significantly larger than the other comparisons (paired t-test, p<0.001). If the neural trajectories are constrained in the 10D space, the Other feedback flow field comparisons should have low flow difference (similar to that for the Fixed feedback comparison) and high flow field overlap (similar to that for the Fixed feedback and Time-reversed comparisons). Taken together, these results indicate that neural flow fields and the resulting neural trajectories are highly consistent in all dimensions, regardless of the visual feedback provided to the animal.

**Extended Data Figure 4.**
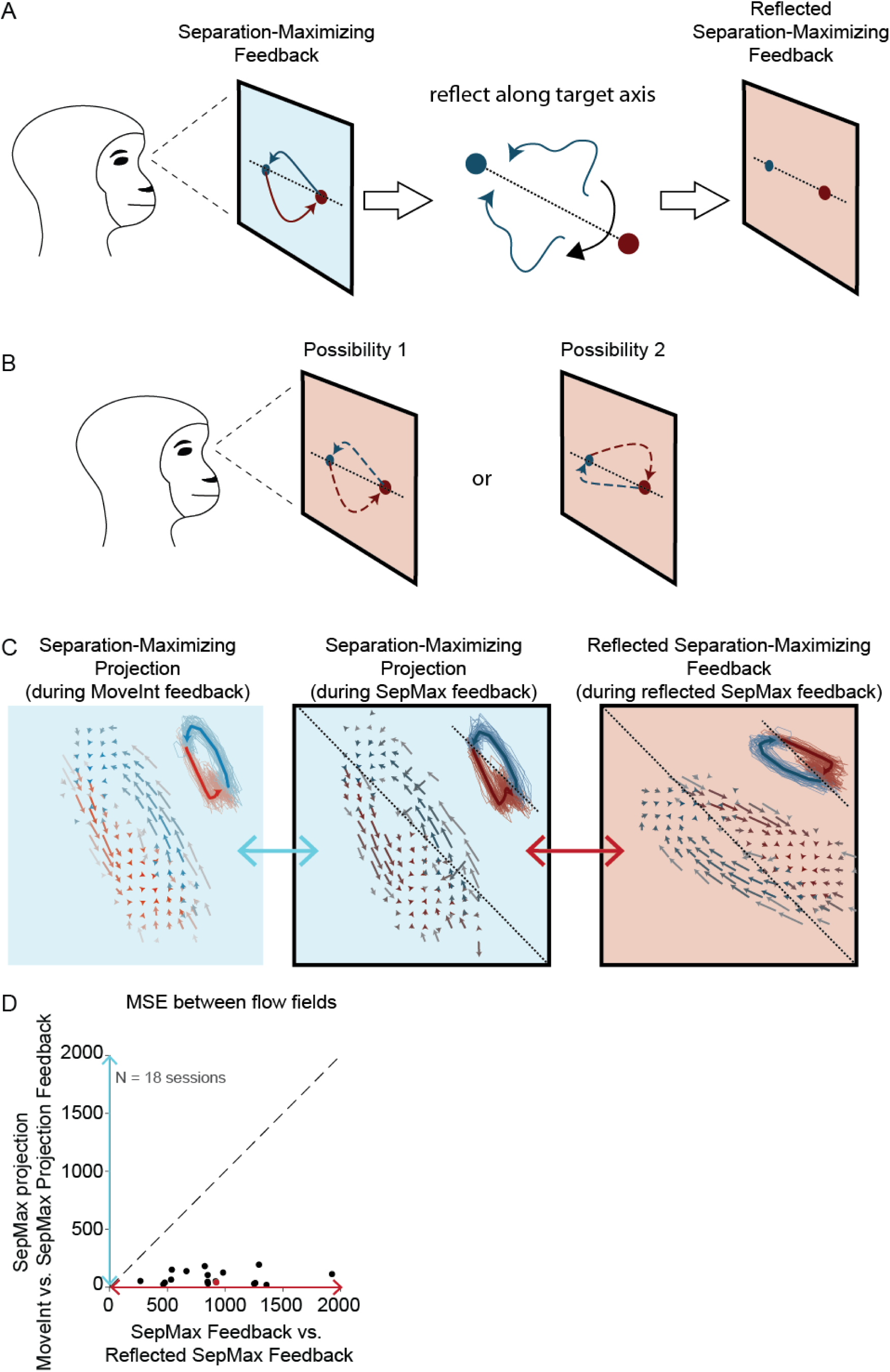
Temporal structure is robust to reflection of the workspace. We assessed if the activity time courses indicate underlying network constraints or a visual preference of the monkey. A. The SepMax projection is unique up to a reflection about the target axis (see Methods). In a separate set of 18 experiments, we presented the animal with both reflections of the matrix *O* (Eqn. 13). To do this, we reflected the orientation of the identified SepMax projection about the target axis to produce a “reflected SepMax” mapping. We then provided the reflected SepMax mapping as feedback to the monkey. B. If the observed temporal structure arose from a visual preference of the monkey, the trajectories under the reflected SepMax feedback would continue to show the structure observed in the SepMax projection (possibility 1). However, if the trajectories arose from underlying network constraints, the trajectories under the reflected SepMax feedback would also be reflected (relative to the SepMax trajectories; possibility 2). C. Flow fields from an example session during BCI control using both SepMax and reflected SepMax mappings. Cursor trajectories are shown as insets. The SepMax projection was identified from the neural activity generated while the animal was receiving visual feedback in the MoveInt projection (left). The animal was provided visual feedback of the SepMax projection (center) and the reflected SepMax projection (right). We observed that the orientation of the cursor trajectories under the reflected SepMax feedback were reflected relative to the trajectories under the SepMax feedback, consistent with possibility 2. D. The flow fields indicated that the trajectory curvature arises from underlying network constraints rather than the animal’s visual preference. We calculated the difference between the flow fields (see Methods) in the SepMax projection and the reflected-SepMax projection. As a benchmark for similar flow fields, we compared this to the difference between the flow fields in the SepMax projection during MoveInt and SepMax feedback. The MSE between the flow fields in the SepMax and reflected SepMax projections (horizontal axis) was significantly larger than the MSE between the flow fields of the SepMax projection in the MoveInt and SepMax feedback conditions (vertical axis; 18/18 sessions Wilcoxon rank sum test, p<0.001). The example session shown in panel (C) is indicated by the red dot in (D).

**Extended Data Figure 5.**
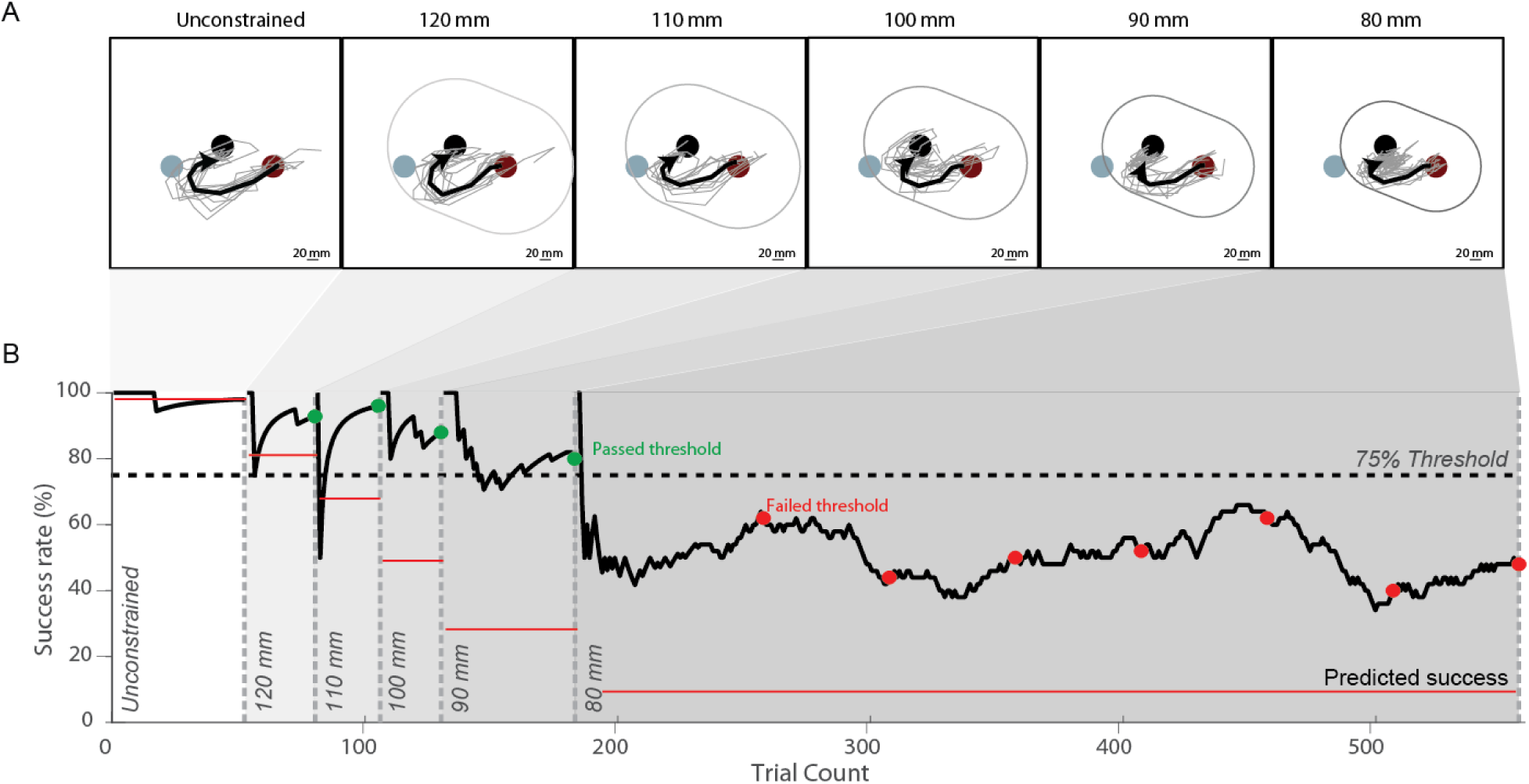
Example of an instructed path experimental session. Animals were instructed to move their BCI cursor from the red target to the black “intermediate” target while keeping the cursor within the visual boundary. For reference, the blue circle indicates the location of the other target in the two-target task, but was not shown to the animal in this task. To encourage the animals to modify their trajectories, we gradually decreased the size of the boundary diameter over the course of each experiment. A. Cursor trajectories for individual trials (thin traces) and averaged across trials (thick traces) to the intermediate target (black circle) during unconstrained trials (far left panel) and in the presence of visual boundaries of decreasing diameters (panels from left to right). As the size of the boundary is reduced, the qualitative structure of the trajectories does not change. B. Success rate over the course of an instructed path experiment. Every 25 trials we checked to see if the success rate was greater than the pre-determined threshold (dashed line). If so (green dots), the boundary was reduced in size. If the animal failed to meet the success rate threshold (red dots), the size was maintained for an additional block of 25 or 50 trials (see Methods). This procedure was continued for a minimum of 500 trials. We compared the observed (thick black) success rate in response to the visual boundary to the predicted (thin red) success rate, computed by applying the same boundaries to the unconstrained trial trajectories. The observed success rate was greater than the predicted success rate, indicating that the animal was responding to the boundary but was unable to change the initial angle of the cursor trajectory (thick traces in panel A).

**Extended Data Figure 6.**
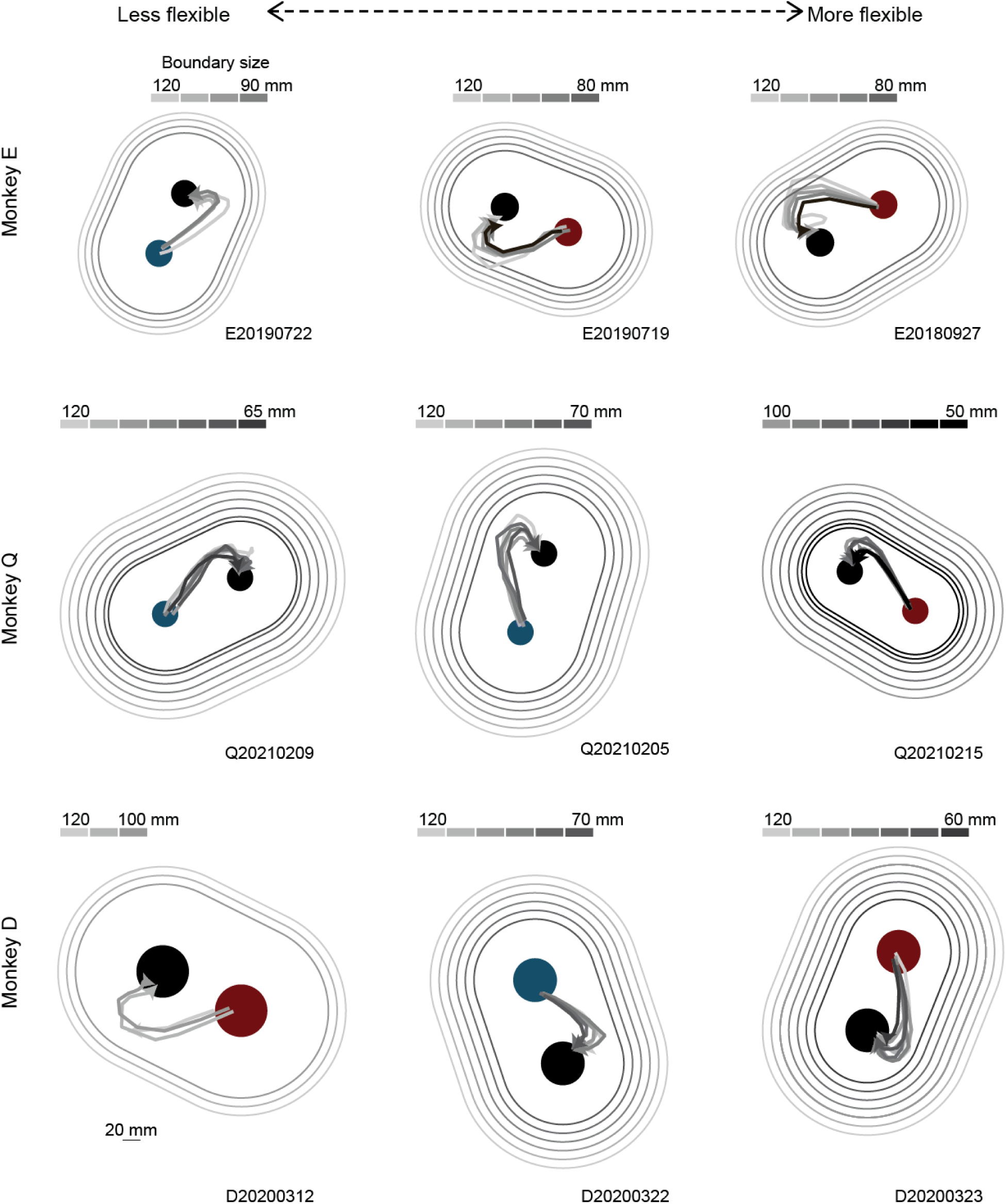
There is minimal flexibility in the cursor trajectory when monkeys are directly challenged to follow an instructed path. Here we show trial-averaged cursor trajectories during instructed path trials for the same sessions as in Extended Data Fig. 2. Sessions are ordered from less flexible (left) to more flexible (right). Flexibility is quantified using the initial angle metric (see Methods). Starting target locations are shown in red or blue, and intermediate targets are shown in black. Trial-average cursor trajectories are plotted for each boundary size (same convention as Fig. 7C). The change in the initial angle is minimal over the course of an instructed path experiment, even in the “more flexible” sessions (cf. Fig. 7). The initial angle does not approach the direct path to the intermediate target.

**Extended Data Figure 7.**
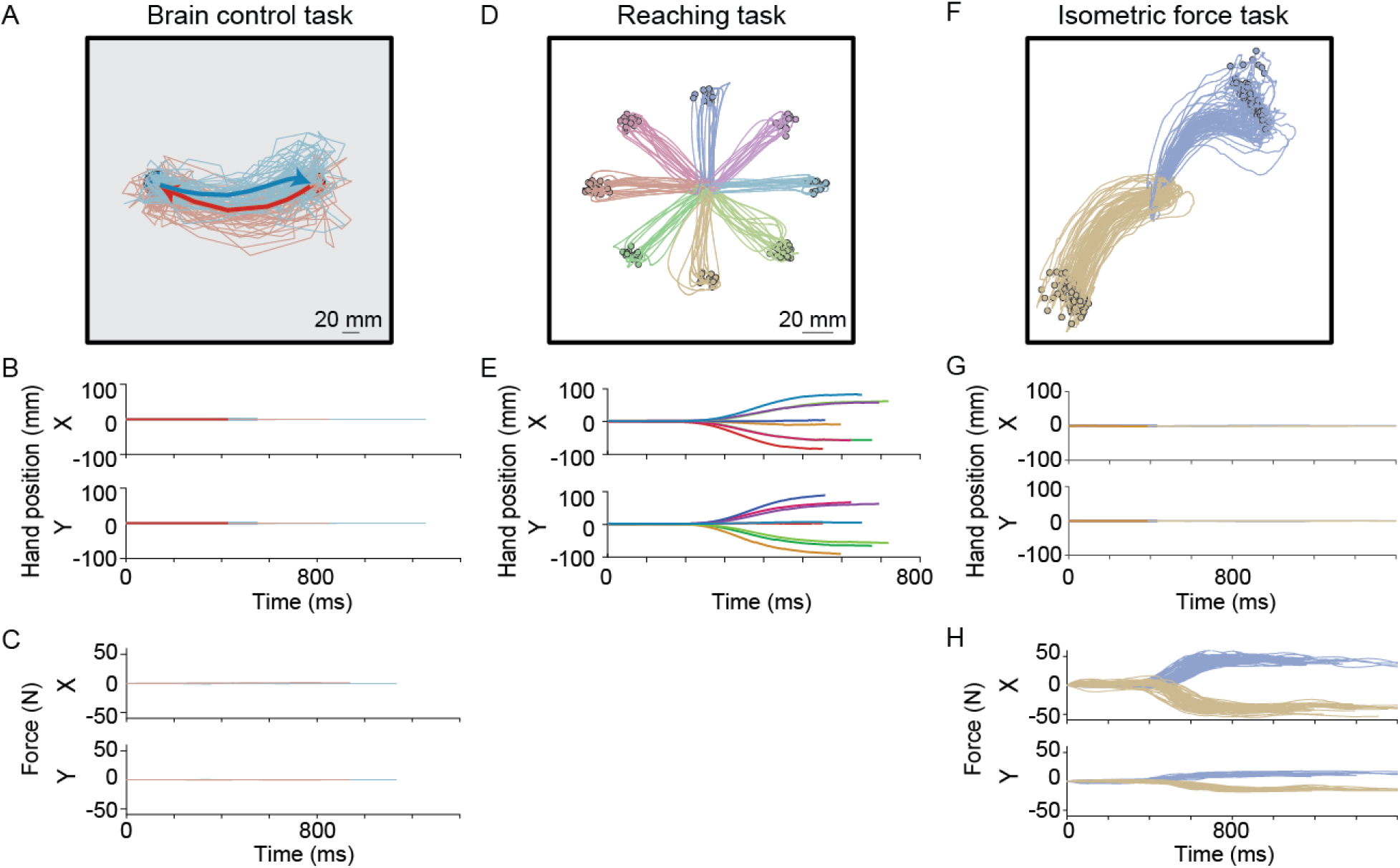
. Arm movements are minimal during BCI cursor control. A. Example cursor trajectories during a two-target BCI task. B. Hand position as a function of time for the two-target BCI trials shown in (A). C. Force produced at the touch bar during the trials shown in (A). D. Example of hand positions during a center-out reaching task. E. Hand position as a function of time for the center-out reaching task trials in (D). Note that changes in hand position during BCI trials (B) is substantially smaller than that observed during center-out reaching trials (E). F. Example cursor trajectories during an isometric force task. For the isometric force task, the monkey applied force to the touch bar. We mapped the exerted force to cursor kinematics, allowing the monkey to acquire force targets. G. Hand position as a function of time for the isometric force task trials shown in (F). H. Force as a function of time for the isometric force task trials shown in (F). Forces exerted on the force bar during BCI control (C) were negligible when compared to those exerted on the force bar during the isometric force task (H).

**Extended Data Figure 8.**
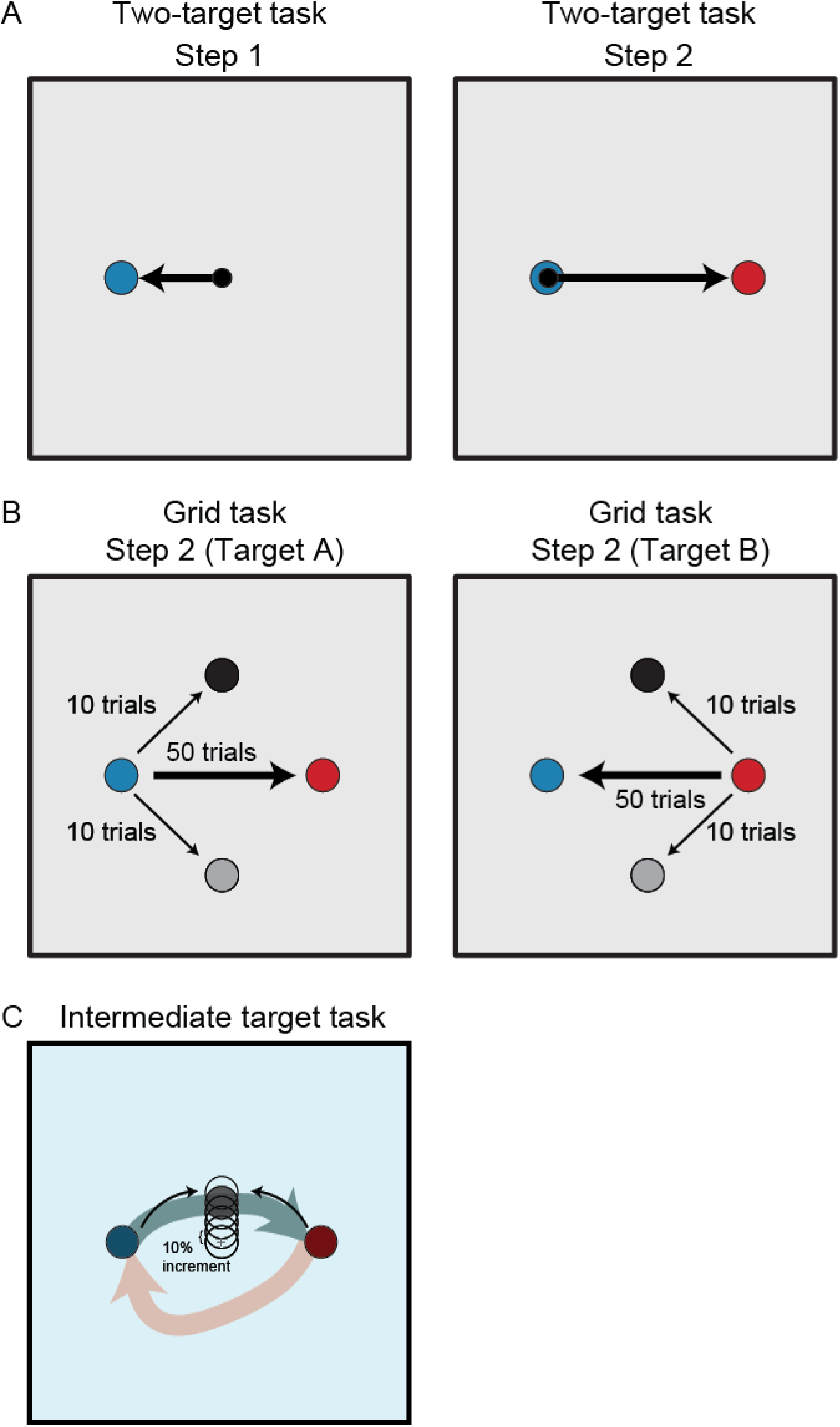
Two-target task, grid task and intermediate target task. There are four possible target pairs. The examples illustrated for each task are for the left-right target pair. A. Two-target task. The two-target task consists of two steps. In step 1 (left), the monkey moved the cursor (small black circle) to a peripheral target (blue circle). Upon completing step 1, step 2 (right) ensued. The monkey moved the cursor to the diametrically opposed target (red circle). B. Grid task. In each session, the same peripheral target pair that was used for the two-target task was used in the grid task. Step 1 for the grid task is the same as that for the two-target task (panel A, left). For step 2, there were three possible target locations: the diametrically opposed target (blue or red circles) or two targets perpendicular to the main target pair (black and gray circles). The probabilities of the targets were weighted so that for a given start target there were 50 trials to the diametrically opposite target and a 10 trials to each of the other two targets. C. Determination of target position for the intermediate target task. To determine the intermediate target position, the monkey first acquired target A or target B (blue and red circles), and then was presented with an intermediate target (open black circles). The location of the intermediate target started at the center of the workspace (gray ‘+’) and we gradually increased the distance from the center in 10% increments of the distance to the peripheral target (open black circles) until the success rate began to decline. Then, we slightly reduced the distance to ensure that the final position of the intermediate target (shaded black circle) was as close as possible to the path defined by blue arrow, while still reachable from the red start target. This iterative approach was necessary to ensure that the intermediate target could be acquired from both start targets.

**Extended Data Figure 9.**
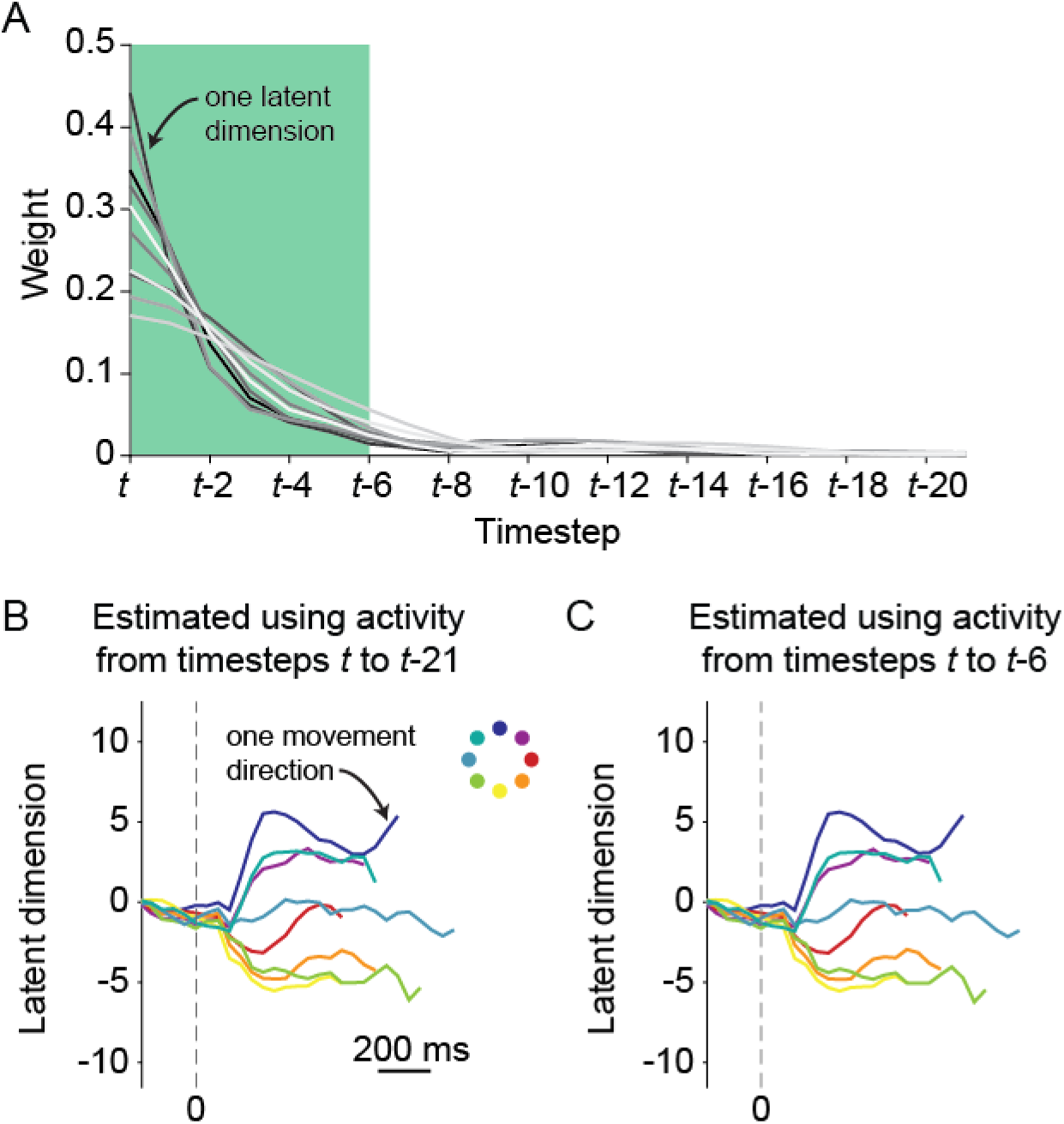
Characterizing GPFA smoothing. A. The latent state estimate at time point *t* (***ẑ***_*t*_) is formed by taking a weighted linear combination of the neural activity at the current and previous time points, where the weights are defined by the smoothing matrix *M* (Eqn. 5). Here we analyze how neural activity at each time point contributes to ***ẑ***_*t*_ by examining the weights in one row of *M*, which corresponds to one latent dimension (one trace). The most recent time bins have the greatest contribution to ***ẑ***_*t*_, whereas the time bins farther into the past contribute less to ***ẑ***_*t*_. We defined *M* based on 22 time steps and used only the weights corresponding to time steps *t* to *t* − 6 only (green shaded area). We truncated the contribution from time steps beyond t-6 for two reasons: 1) so that neural activity at the end of one trial would not influence the estimated latent states at the start of the following trial, and 2) for computational efficiency. B, C. Example single-trial neural trajectories along one latent dimension. The neural trajectories estimated using all 22 time steps of neural activity (*t* to *t* − 21, i.e., no truncation; panel B) were similar to those estimated using only the 7 most recent time steps of neural activity (*t* to *t* − 6, i.e., with truncation; panel C). Each colored trace corresponds to one movement direction. The trajectories in (B) and (C) look similar because the weights for time points beyond *t* − 6 are small (as shown in A).

**Extended Data Figure 10.**
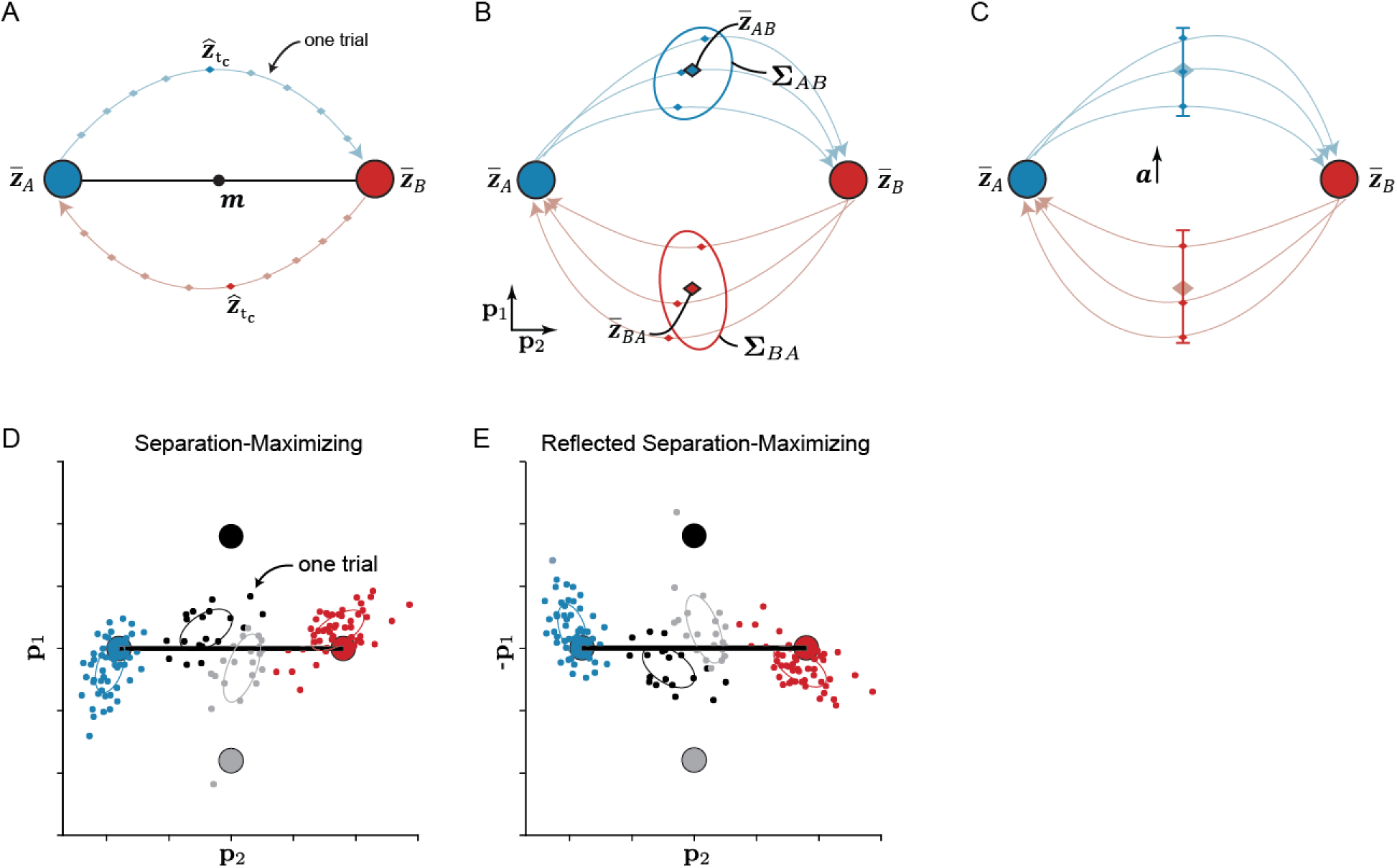
Identifying the SepMax projection. A. The SepMax mapping was designed to highlight projections of neural activity in which A-to-B neural trajectories are maximally separated from B-to-A neural trajectories. The first step in identifying the projection was to find the midpoint of each neural trajectory. For each trial, we defined the midpoint of the neural trajectory, ***ẑ***_*tc*_ ∈ *R*^10×1^, to be the time point whose projection is closest to the midpoint, ***m*** ∈ *R*^10×1^, between the starting points of the neural trajectories ***z̄***_*A*_ and ***z̄***_*B*_. The vectors ***z̄***_*A*_ and ***z̄***_*B*_ ∈ *R*^10×1^ are the trial-averaged starting locations for the A-to-B and B-to-A trajectories during the two-target task. Time points are indicated as dots along the trajectory. B. Conceptual illustration of the features that define the objective function (Eqn. 10) used to identify the SepMax mapping. ***z̄***_*AB*_ and ***z̄***_*BA*_ ∈ *R*^10×1^ are the trial-averaged midpoints of the A-to-B and B-to-A trajectories, and ∑_*AB*_ and ∑_*BA*_ ∈ *R*^10×10^ are the covariance matrices describing the trial-to-trial scatter of the midpoints of the A-to-B and B-to-A trajectories, respectively. C. The discriminability index (*d*′) used to measure how distinct the neural trajectories are between the A-to-B and B-to-A conditions. We defined an axis, ***a***, separating the trial-averaged midpoints of the two conditions (i.e., ***z̄***_*AB*_ and ***z̄***_*BA*_). We projected the midpoints of the A-to-B trajectories (blue) and the B-to-A trajectories (red) onto ***a***. Using the means and variances of these projections, we calculated *d*′ (Eqn. 15). D, E. Choosing the orientation (i.e., matrix *O*; Eqn. 13) of the SepMax projection. The SepMax projection is determined up to a reflection about the target axis (black line). Panels D and E represent two candidate SepMax mappings. We chose the orientation of the SepMax projection based on visual inspection of the endpoints of the trajectories during the grid task. Specifically, the SepMax projection was chosen such that the endpoints of the neural trajectories to the orthogonal grid targets (small black and gray dots) were closest to the associated target location (large black and gray circles). In this example, the mapping shown in (D) would be selected as the SepMax mapping because the small black and gray dots appear on the same side of the target axis as the black and gray targets, respectively. The mapping shown in (E) would be chosen as the reflected SepMax projection (Extended Data Fig. 4). Note that the small black and gray dots appear on the opposite side of the target axis as the black and gray targets, respectively. This color convention differs from the convention throughout the rest of the manuscript in which trajectories are colored by the start target.

**Supplemental Table 1.** Instructed path task experiment details for Monkeys E, D and Q. Targets are arranged around a circle at 45 ° intervals. The start target angle indicates the position of the start target. The intermediate target is placed at the indicated angle (°) and distance from the center of the workspace (mm). The range of the boundary size tested for each experiment is reported as the largest and smallest boundary (mm).

|  |  |  | Intermediate target |  | Boundary |  |
| --- | --- | --- | --- | --- | --- | --- |
| Monkey | Session | Start target angle (°) | Angle (°) | Distance from center (mm) | Size of largest (mm) | Size of smallest (mm) |
| E | 20180924 | 225 | 135 | 45 | 120 | 70 |
|  | 20180925 | 315 | 225 | 45 | 120 | 80 |
|  | 20180926 | 90 | 180 | 54 | 120 | 90 |
|  | 20180927 | 0 | 270 | 45 | 120 | 80 |
|  | 20181015 | 315 | 45 | 27 | 100 | 80 |
|  | 20181016 | 180 | 90 | 36 | 80 | 60 |
|  | 20181017 | 90 | 0 | 27 | 110 | 90 |
|  | 20181018 | 45 | 315 | 27 | 110 | 80 |
|  | 20181019 | 0 | 90 | 27 | 110 | 60 |
|  | 20181021 | 135 | 225 | 9 | 110 | 30 |
|  | 20181022 | 270 | 180 | 45 | 110 | 110 |
|  | 20181023 | 225 | 315 | 27 | 110 | 70 |
|  | 20181024 | 315 | 45 | 36 | 110 | 100 |
|  | 20181025 | 90 | 180 | 18 | 110 | 50 |
|  | 20181026 | 45 | 315 | 27 | 110 | 60 |
|  | 20181028 | 180 | 90 | 45 | 110 | 40 |
|  | 20181029 | 315 | 45 | 36 | 110 | 110 |
|  | 20181031 | 0 | 90 | 27 | 120 | 110 |
|  | 20181101 | 90 | 180 | 45 | 110 | 110 |
|  | 20181107 | 180 | 270 | 36 | 110 | 90 |
|  | 20190201 | 0 | 270 | 36 | 120 | 120 |
|  | 20190204 | 45 | 135 | 36 | 120 | 80 |
|  | 20190207 | 90 | 0 | 27 | 120 | 70 |
|  | 20190208 | 315 | 45 | 45 | 120 | 90 |
|  | 20190719 | 0 | 90 | 36 | 120 | 80 |
|  | 20190722 | 225 | 135 | 36 | 120 | 90 |
|  | 20190723 | 90 | 180 | 27 | 120 | 60 |
|  | 20190724 | 135 | 45 | 27 | 120 | 60 |
| D | 20200312 | 0 | 90 | 45 | 120 | 100 |
|  | 20200313 | 180 | 270 | 36 | 150 | 130 |
|  | 20200316 | 135 | 225 | 36 | 120 | 50 |
|  | 20200318 | 45 | 315 | 45 | 120 | 70 |
|  | 20200319 | 90 | 0 | 18 | 120 | 100 |
|  | 20200320 | 0 | 270 | 36 | 120 | 110 |
|  | 20200321 | 180 | 90 | 9 | 120 | 40 |
|  | 20200322 | 135 | 225 | 45 | 120 | 70 |
|  | 20200323 | 90 | 180 | 36 | 120 | 60 |
| Q | 20210204 | 225 | 270 | 26.1 | 120 | 70 |
|  | 20210205 | 270 | 0 | 27 | 120 | 70 |
|  | 20210207 | 315 | 225 | 27 | 120 | 65 |
|  | 20210208 | 180 | 0 | 0 | 120 | 70 |
|  | 20210209 | 225 | 315 | 31.5 | 120 | 65 |
|  | 20210211 | 315 | 225 | 22.5 | 100 | 40 |
|  | 20210213 | 225 | 315 | 34 | 85 | 70 |
|  | 20210215 | 315 | 225 | 22.5 | 100 | 50 |
|  | 20210217 | 315 | 225 | 17 | 100 | 40 |
|  | 20210218 | 180 | 90 | 17 | 90 | 50 |
|  | 20210220 | 270 | 180 | 21.25 | 110 | 45 |
|  | 20210221 | 0 | 90 | 25.5 | 120 | 55 |
|  | 20210222 | 270 | 180 | 12.75 | 100 | 55 |

